# A cryo-EM structure of metazoan TRAPPIII, the multisubunit complex that activates the GTPase Rab1

**DOI:** 10.1101/2020.12.17.423307

**Authors:** Antonio Galindo, Vicente J. Planelles-Herrero, Gianluca Degliesposti, Sean Munro

## Abstract

The TRAPP complexes are highly conserved nucleotide exchange factors, with TRAPPIII activating Rab1 and TRAPPII acting primarily on Rab11. The two complexes share a core of small subunits that affect nucleotide exchange, but are distinguished by additional large subunits that are essential for activity in vivo, and are mutated in a range of human disorders. Crystal structures of the core subunits have revealed the mechanism of Rab activation, but how and why the large subunits associate with the core remains unclear. We report here a cryo-EM structure of the entire TRAPPIII complex from *Drosophila*. The TRAPPIII-specific subunits TRAPPC8 and TRAPPC11 hold the catalytic core like a pair of tongs, with TRAPPC12 and TRAPPC13 positioned at the joint between them. TRAPPC2 and TRAPPC2L link the core to the two large arms, with the interfaces containing residues affected by disease-causing mutations. The TRAPPC8 arm is positioned such that it would contact bound Rab1, indicating how the arms could alter the Rab specificity of the core. A lower resolution structure of TRAPPII shows a similar architecture, and suggests that the TRAPP complexes evolved from a single ur-TRAPP.

## Introduction

Small GTPases of the Rab family are major regulators of membrane traffic and organelle location in eukaryotic cells. Upon activation, they recruit to specific membranes a diverse set of effectors including molecular motors, tethering factors, and regulators of both GTPases and phosphoinositides. The internal organisation of the cell thus depends on these GTPases being activated only in the correct location. This activation is mediated by nucleotide exchange factors (GEFs) that bind the inactive GDP-bound form and catalyse the release of GDP and replacement with GTP. It has become clear that the primary determinant of the spatial accuracy of GTPase activation is the location of the relevant GEFs, and hence understanding their structure and regulation is key to understanding the organisation of the cell (Barr, 2013; Blümer et al., 2013). The Transport Protein Particle (TRAPP) GEFs were discovered in yeast, and subsequently found to be conserved in all known eukaryotes (Brunet and Sacher, 2014; Kim et al., 2016; Klinger et al., 2013; Sacher et al., 1998). In most species examined to date, including metazoans, there are two versions, TRAPPII and TRAPPIII, with TRAPPI now thought to be a subcomplex that appears in vitro during isolation of the other two (Brunet et al., 2012; Choi et al., 2011; Thomas et al., 2017). TRAPPIII activates Rab1, a master regulator of both the early secretory pathway and autophagy, whilst TRAPPII primarily activates Rab11, an essential player at the late Golgi where it acts in recycling from endosomes, and traffic to the surface (Cai et al., 2005; Jones et al., 2000; Wang et al., 2000). Rab1 and Rab11 are two of the five members of the Rab family that are present in all eukaryotes, and both are essential for the viability of all organisms so far examined (Diekmann et al., 2011; Kloepper et al., 2012). TRAPPII also has some activity on Rab1, although the in vivo significance of this is unresolved (Ke et al., 2020; Thomas and Fromme, 2016; Yamasaki et al., 2009). The TRAPP complexes have also been proposed to have additional roles in various processes including tethering of COPII vesicles, meiotic cytokinesis, ciliogenesis, and lipid droplet homeostasis (Cai et al., 2007; Li et al., 2017; Robinett et al., 2009; Westlake et al., 2011). Consistent with the TRAPP complexes acting in key cellular processes, mutations in many of their subunits have been found in a range of familial conditions or “TRAPPopathies”, including neurodevelopmental disorders, muscular dystrophies and skeletal dysplasias (Gedeon et al., 1999; Matalonga et al., 2017; Sacher et al., 2019).

The two TRAPP complexes share a core of seven small subunits, one of which is present in two copies to make an octamer (Figure 1A). This core is sufficient to activate Rab1 in vitro (Kim et al., 2006; Riedel et al., 2017). In most species, TRAPPIII has four additional unique subunits, TRAPPC8, TRAPPC11, TRAPPC12 and TRAPPC13, with the former two being essential in metazoans for cell viability and Rab1 recruitment, indicating that the core is not sufficient to correctly activate Rab1 in vivo (Lamb et al., 2016)(Kim et al., 2016; Wendler et al., 2010) (Riedel et al., 2017). TRAPPII has two additional unique subunits, TRAPPC9 and TRAPPC10, that are required for Rab11 activation both in vitro and in vivo (Riedel et al., 2017; Thomas et al., 2017). Crystallographic studies of the individual core subunits and their subcomplexes have revealed that the centre of the core comprises two longin domain proteins, TRAPPC1 and TRAPPC4, consistent with longin domains being present in several other Rab GEFs (Cai et al., 2008; Kim et al., 2006; Levine et al., 2013). Flanking these are TRAPPC5, TRAPPC6, and two copies of TRAPPC3, all of which fold into a distinct TRAPP domain that appears to have emerged in archaea, consistent with TRAPP being a universal feature of eukaryotes (Kümmel et al., 2005; Zaremba-Niedzwiedzka et al., 2017). Rab1 binds to this core region, with a subcomplex of TRAPPC1, TRAPPC3 and TRAPPC4 being sufficient for GEF activity in vitro (Cai et al., 2008; Kim et al., 2006). Finally, TRAPPC2 and TRAPPC2L, two longin-like proteins, are found at the two ends of the core and are required to connect the core to the specific subunits of TRAPPII and TRAPPIII (Montpetit and Conibear, 2009; Zong et al., 2011).

**Figure 1.**
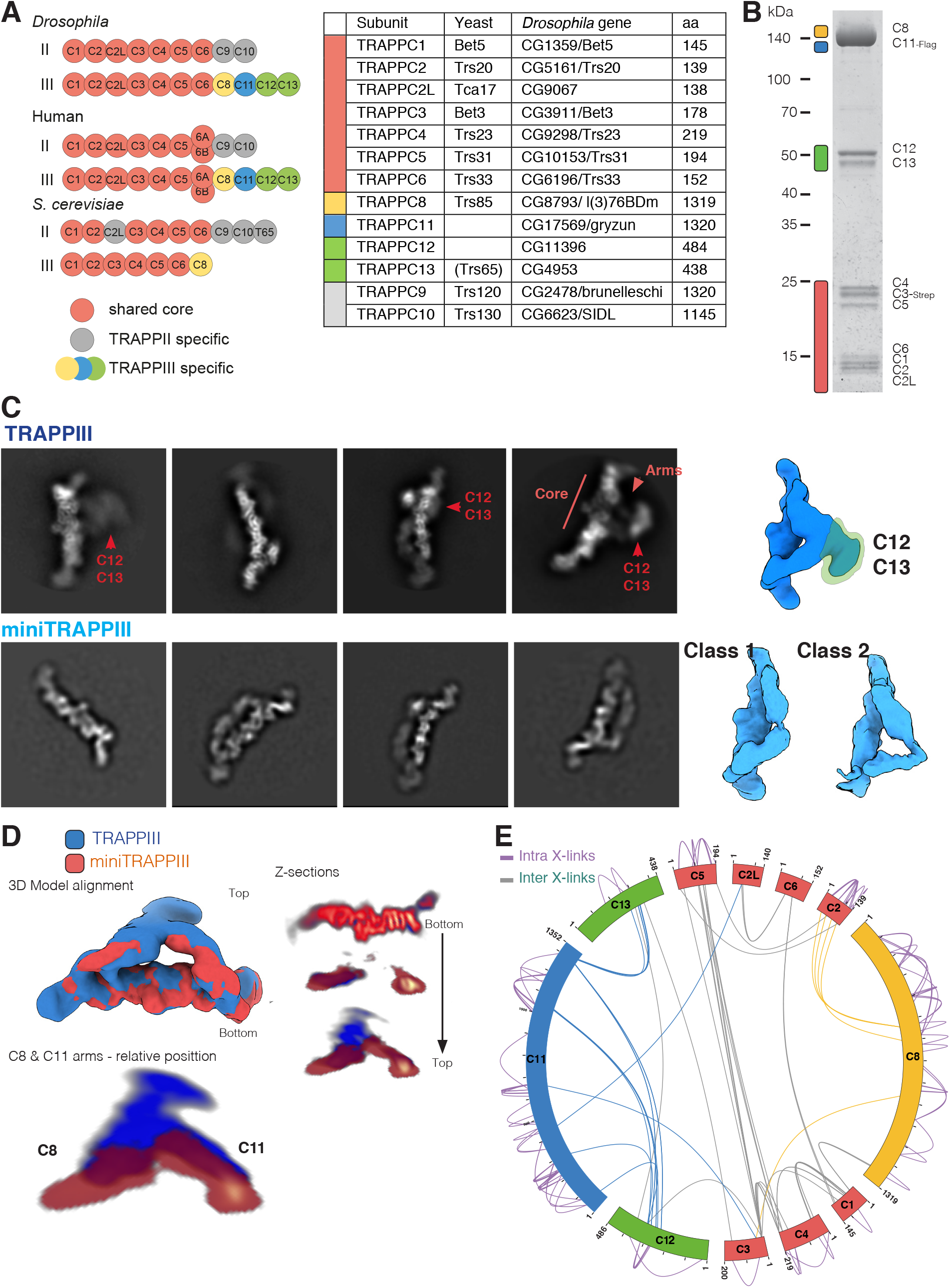
Single particle imaging of the *Drosophila* TRAPPIII complex. (A) Subunit structure of the TRAPP complexes from *Drosophila*, humans and *S. cerevisiae*. The *S. cerevisiae* TRAPPII subunit Trs65 is distantly related to TRAPPC13, but appears to a paralogue of TRAPPC13 that arose in fungi, rather than a true orthologue. (B) Coomassie blue-stained gel of recombinant *Drosophila* TRAPPIII complex. The subunits of TRAPPIII are indicated. (C) Right: representative 2D class averages of TRAPPIII and miniTRAPPIII. Left: Low resolution 3D models of TRAPPIII and miniTRAPPIII. The density corresponding to TRAPPC12 and TRAPPC13 missing in miniTRAPPIII is indicated on TRAPPIII. (D) Alignment of 3D models of TRAPPIII and miniTRAPPIII. Representative Z-sections of the alignment are shown. Maximum correlation is found at the core region (bottom). A top plane from the Z-sections is enlarged. Density missing in the absence of TRAPPC12 and TRAPPC13 is apparent, with the arms also moving towards the core. (E) Circos-XL plots showing the distribution of all DSBU cross-links for the whole TRAPPIII complex. Each protein is represented as a coloured segment (core subunits: red, specific subunits: TRAPPC8 as yellow, TRAPPC11 as blue, TRAPPC12 and TRAPPC13 as green), with the amino acid residues indicated on the outside of the plot. The relative position of the cross-link represents its location within the primary sequence. Inter-molecular cross-links are depicted as purples lines on the outside of the plot and intra-molecular cross-links as green lines inside of the plot.

The architectures of the entire TRAPP complexes are less well understood. Low resolution images of budding yeast TRAPPIII obtained with negative stain EM show that Trs85, the orthologue of TRAPPC8, is attached to one end of the core via the orthologue of TRAPPC2 (Tan et al., 2013). However, fungal Trs85 represents just the N-terminal half of TRAPPC8 and it lacks the C-terminal 600-700 residues present in plants and metazoans. In addition, *S. cerevisiae* is distinct from most other species including many other fungi, in that it lacks the TRAPPIII subunits TRAPPC11, TRAPPC12 and TRAPPC13, even though the former is essential in both *Drosophila* and mammals, and so its TRAPPIII is simpler (Figure 1A) (Kalde et al., 2019; Kim et al., 2016; Riedel et al., 2017; Rosquete et al., 2019; Wendler et al., 2010). Negative stain EM images of yeast TRAPPII show that the entire complex comprises Trs120 (TRAPPC9) and Trs130 (TRAPPC10) flanking the core, with this structure then dimerising with a second copy via Trs65 that links together the ends of TRAPPC9 in one copy to TRAPPC10 in the other (Pinar et al., 2019; Yip et al., 2010). However, Trs65 seems to have evolved in the yeast lineage as a distinct variant of TRAPPC13, with some other fungi having both proteins, suggesting that this form of TRAPPII is unique to a subset of fungi (Pinar et al., 2019).

To obtain insight into the architecture of the metazoan TRAPP complexes we expressed recombinant TRAPPII and TRAPPIII using the *Drosophila* subunits. Single particle cryo-EM was used to obtain a structure of the TRAPPIII. This structure resolves the uncertainty about the organisation of the subunits of the core, shows how all of the additional subunits are arranged in the complex, maps the interfaces between the core and these subunits, including residues involved in genetic disease, and reveals how these additional subunits could regulate Rab binding and hence allow the core to act on different GTPases in the two different complexes.

## Results

### Biochemical characterisation of the metazoan TRAPP complexes

In previous work we developed a protocol to express and purify recombinant forms of the *Drosophila* TRAPP complexes (Figure 1B)(Riedel et al., 2017). We reported that the purified complexes are functional, with both TRAPPII and TRAPPIII having nucleotide exchange activity towards Rab1, while only TRAPPII has detectable exchange activity on Rab11. Further characterisation of these complexes shows that they are both monodisperse and monomeric, as indicated by both multi-angle light scattering coupled with size exclusion chromatography (SEC-MALS) and interferometric scattering microscopy (iSCAT) (Figure S1A-S1D).

The two largest subunits of TRAPPIII, TRAPPC8 and TRAPPC11, are essential for viability of mammalian cells, and Trs85, the yeast orthologue of TRAPPC8 has been shown to bind directly to the core via TRAPPC2 (Brunet et al., 2013; Tan et al., 2013; Taussig et al., 2014). However, TRAPPIII has two further subunits TRAPPC12 and TRAPPC13 whose location in the complex is unknown. Expressing TRAPPIII without the TRAPPC12 and TRAPPC13 subunits reduces its size by ~100 kDa, close to the combined weight of two subunits, indicating that their absence does not affect the binding of the other nine subunits (Figures S1A-S1D). This “miniTRAPPIII” complex is still able to activate Rab1 but, like the complete complex, it has no detectable activity on Rab11 (Figure S1E).

### Cryo-EM analysis of the TRAPPIII complex

To obtain new insights into the architecture of TRAPPIII, we applied electron microscopy (EM) to examine its structure. When examined by negative staining, TRAPPIIII particles appeared homogeneous in overall size, and of rod-like or triangular appearance (Figure S2A). The particles appeared similar in cryo-EM micrographs, and 2D class averages showed clear elements of secondary structure (Figure S2B). However, initial attempts to produce a reliable 3D reconstruction failed, and we noticed that several 2D class averages had a threefold symmetry, forming an equilateral triangle (Figure S2B). This seemed inconsistent with TRAPPIII being a monomer rather than a trimer in solution, suggesting that these symmetrical particles were due to overfitting of 2D projections. Moreover, in the case of the miniTRAPPIII, similar narrow rod and triangular particles were found in the cryo-EM micrographs, but threefold symmetrical particles rarely appeared among the 2D class averages (Figure S2C). We could identify two main 2D classes of rod-like particles, one of them that partially resembled the low resolution structure of yeast TRAPPIII obtained by negative stain (Tan et al., 2013), and another similar to it but with an additional density on one of its edges. We used a 3D reconstruction map of the latter as a reference map for a 3D classification of the TRAPPIII class 2D averages. This classification resulted in three different classes. The one with the best resolution and highest number of particles corresponded to an irregular triangular shape which resembled that which we observed with negative stain (Figures 1C and S2A)

We reanalysed cryo-EM images of miniTRAPPIII following a similar strategy to that used for TRAPPIII. The comparison between the TRAPPIII and miniTRAPPIII class 2D averages and 3D models revealed that the flat rod was similar, but in miniTRAPPIII there was density missing at the region of interaction between the two arms (Figures 1C and 1D). We concluded TRAPPC12 and TRAPPC13 were located in this region of TRAPPIII, forming one of the vertexes of the triangle (Figure 1C)

### Refinement of TRAPPIII density map

Following averaging and refinement, the initial density map of TRAPPIII comprises an elongated flat rod with a small protrusion in the middle, and two arms that are attached at the two ends of this rod (Figure 1C). Secondary structure was better resolved inside the flat rod rather than in the two arms, which indicated two problems. Firstly, there is incomplete angular distribution of the particles due to a preferred orientation of the complex on frozen grids, (Figure S3). Secondly, the arms attached to the rod are somewhat flexible. To address the first problem, we imaged grids tilted by 19°, and combining the tilted and non-tilted datasets gave a 3D reconstruction with a nominal resolution of 5.8 Å (Figure S4A).

After several tests, we divided the reconstructed map into three different bodies for focused refinement. This approach resulted in a 4.27 Å resolution for a body containing the core subunits that form the flat rod, 4.57 Å for a second body containing one arm and the TRAPPC12 and TRAPPC13 subunits, and 5.5 Å for a third body formed by the other arm (Table 1 and Figures S4A-S4B).

**Table 1.** Cryo-EM Data Collection, Refinement, and Validation Statistics.

|  | TRAPIII | miniTRAPIII |
| --- | --- | --- |
| **Data collection and processing** |  |  |
| Magnification | 75000x | 105000x |
| Voltage (kV) | 300 | 300 |
| Electron exposure (e-/Å<sup>2</sup>) | 30 | 45.6 |
| Defocus range (μm) | -2.2 / -4 | -1.5 / -2.5 |
| Pixel size (Å) | 1.04 | 1.09 |
| Movies (no.) | 3671 | 3443 |
| Symmetry imposed | C1 | C1 |
| Initial particle images (no.) | 1601314 | 128478 |
| Final particle images (no.) | 353400 | 486758 |
| Map sharpening *B* factor (Å<sup>2</sup>) | -34.32 | -45.77 |
| Map resolution (Å) | 5.8 | 4 |
| FSC threshold | 0.143 | 0.143 |
| Map resolution range (Å) | 4-9 | 3.5-7.5 |
| EMDB accession code | EMD-12056 | EMD-12063 |
| PDB accession code | 7B6R | 7B7O |

| **Refinement** | **Core** | **C8** | **C11** |
| --- | --- | --- | --- |
| Map sharpening *B* factor (Å<sup>2</sup>) | -39.42 | -45.48 | -124.75 |
| Model resolution (Å) | 4.2 | 4.6 | 5.4 |
| FSC threshold | 0.143 | 0.143 | 0.143 |
| Model composition |  |  |  |
| Chains | 8 | 1 | 1 |
| Nonhydrogen atoms | 10470 | 2205 | 4904 |
| Protein residues | 1290 | 269 | 614 |
| Ligands | 0 | 0 | 0 |
| Protein *B* factors (Å<sup>2</sup>) | 214.4 | 302.4 | 161.8 |
| R.m.s. deviations |  |  |  |
| Bond lengths (Å) | 0.005 | 0.005 | 0.006 |
| Bond angles (°) | 1.090 | 1.018 | 1.030 |
| EMDB accession code | EMD-12052 | EMD-12053 | EMD-12054 |
| PDB accession code | 7B6d | 7B6E | 7B6H |

| **Validation** |  |  |  |
| --- | --- | --- | --- |
| MolProbity score | 2.55 | 2.61 | 2.68 |
| Clashscore | 32.13 | 35.59 | 45.19 |
| Poor rotamers (%) | 0.43 | 0 | 0 |
| Ramachandran plot |  |  |  |
| Favored (%) | 89.72 | 89.06 | 90.07 |
| Allowed (%) | 10.20 | 10.94 | 9.77 |
| Disallowed (%) | 0.08 | 0.00 | 0.17 |

### Arrangement of the subunits within the TRAPPIII complex

To help locate the 12 subunits of TRAPPIII within the density map we used cross-linking mass spectrometry to identify lysine residues in proximity to each other (Figure 1E and Table S1). As expected, there were numerous cross-links between the small subunits of the core. Of the large subunits, TRAPPC8 made several links to the TRAPPC2 subunit that is expected to be at one end of the core, as well as a link to TRAPPC3, whilst TRAPPC11 linked to TRAPPC2L and also to TRAPPC3. Association of TRAPPC8 with TRAPPC2 is consistent with what is known of yeast TRAPPIII from the effect of mutations in the TRAPPC2 orthologue, Trs20 (Brunet et al., 2013; Taussig et al., 2014). The TRAPPC12 and TRAPPC13 subunits primarily crosslinked to the C-terminal region of TRAPPC11, indicating that this part of TRAPPC11 is located in this vertex. Taken together, these results allow an unambiguous placement of the subunits within the TRAPPIII density map in which the core sits between arms formed from TRAPPC8 and TRAPPC11 with TRAPPC12 and TRAPPC13 attached at the opposite vertex. The overall shape of the complex is that of two arched arms connected at one vertex with TRAPPC12 and TRAPPC13, and then spreading apart to hold the core between their other ends like a pair of tongs (Figure 2A).

**Figure 2.**
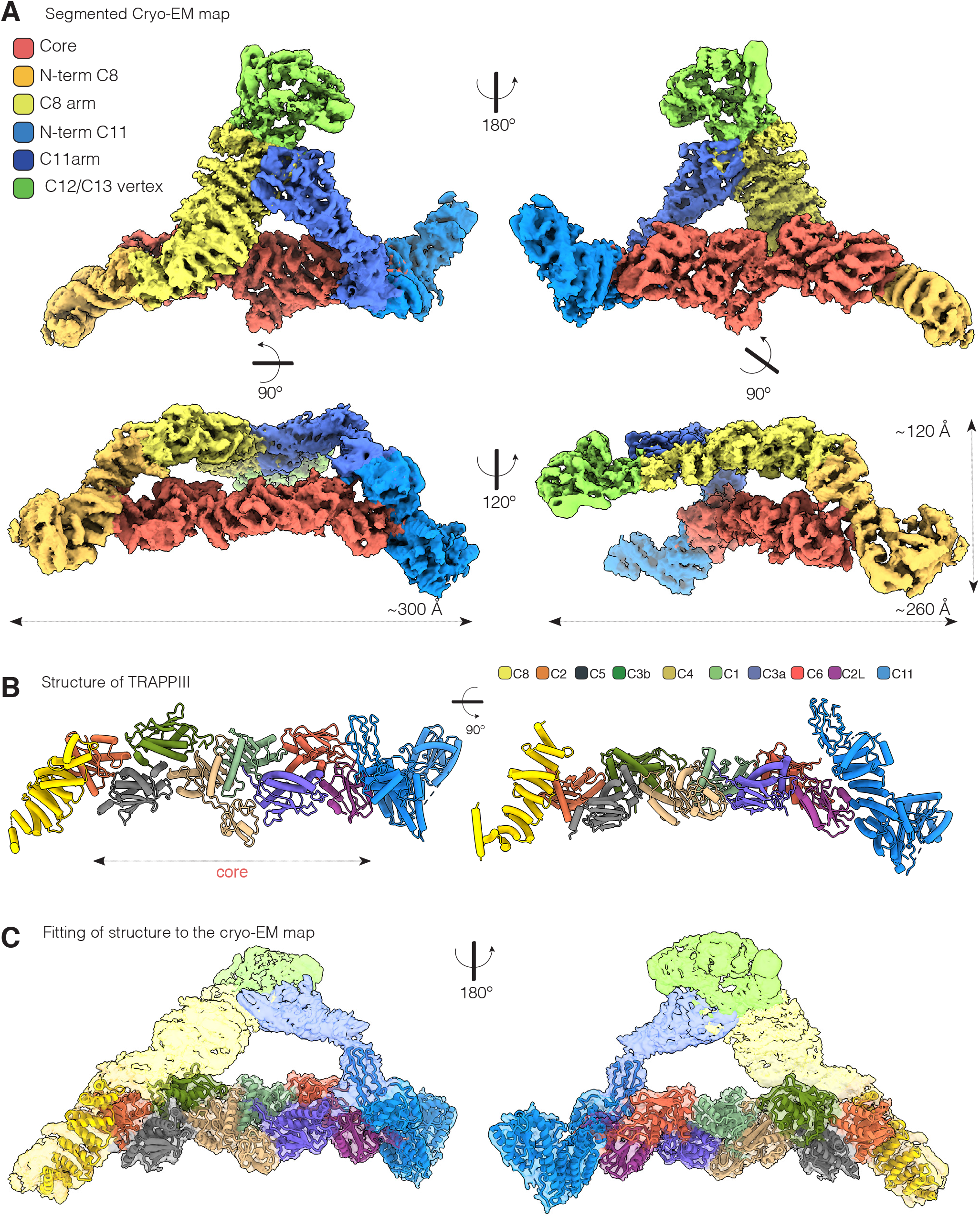
Architecture of the *Drosophila* TRAPP III complexes. (A) Cryo-EM density map coloured according to the location of the TRAPPIII specific subunits: TRAPPC8 (N-terminus: dark yellow; C-terminus: light yellow), TRAPPC11(N-terminus: light blue; C-terminus: dark blue), TRAPPC12 and TRAPPC13 (light green), and the TRAPP Core subcomplex (red). Four different views are shown. (B) Orthogonal views of the partial TRAPPIII structural model. The views are depicted as pipes and planks and coloured by subunits (C1: light green, C2: orange, C2L: magenta, C3a: purple, C3b: dark green, C4: light brown, C5: grey, C6: red, C8: yellow, C11: blue). (C) TRAPPIII structural model as in (B) fitted into the cryo-EM map.

To assign the eight small subunits within the core, we used the crystal structures that have been obtained for several of these subunits from mammals, either singly or in subcomplexes comprising up to four subunits (Jang et al., 2002; Kim et al., 2006; Wang et al., 2014). These were used to model the seven *Drosophila* core subunits which were then built into the density map. The subunits could be readily fitted into the map with the assembly of all seven, with TRAPPC3 being present twice, creating an octamer that forms the central flattened rod (Figures 2B, 2C and S5). As expected, TRAPPC1 and TRAPPC4, which form the catalytic site to activate Rab1 (Cai et al., 2008), are at the centre of the rod, flanked on either side by a TRAPPC3 subunit (C3a and C3b). TRAPPC1 and TRAPPC4 share a longin-domain fold, with TRAPPC4 also having a PDZ-like domain that protrudes from one side. The absence of this domain in TRAPPC1 leaves a groove on the other side of the rod, which is partially occupied by the C-terminal region of one of the two TRAPPC3 subunits (C3b). TRAPPC5 and TRAPPC2 bind one TRAPPC3 (C3b), and TRAPPC6 and TRAPPC2L bind the other (C3a). TRAPPC1, TRAPPC5 and TRAPPC2 are known to be related to TRAPPC4, TRAPPC6 and TRAPPC2L respectively, and so the octamer has an approximate two-fold rotational symmetry, suggesting that it evolved by gene duplications adding, or altering, one half. In the octameric assembly the greatest divergence is between TRAPPC5 and TRAPPC6. TRAPPC5 contains a disordered N-terminal region and an extra C-terminal α-helix that is not present in TRAPPC6 (Figure S5).

### Incorporation of the TRAPP core into the rest of the complex

There are no reported crystal structures for any of the large TRAPP subunits from any species. However, in the regions of TRAPPC8 and TRAPPC11 located on either side of the core the local resolution was suitable for de novo model building. In the case of TRAPPC8, residues 350 to 660 form an α-solenoid of thirteen α-helices that includes the site of interaction with TRAPPC2 (Figure 3). The less well resolved C-terminal region (residues 660-1319) forms the arm that connects to TRAPPC12 and TRAPPC13 (Figure 2C). This part of TRAPPC8 is not present in the yeast orthologue Trs85, consistent with it forming an armless complex (Tan et al., 2013). An α-solenoid is also seen for TRAPPC11, with residues 181-566 forming fifteen α-helices that includes the interaction surface with TRAPPC2L (Figure 4). This region of TRAPPC11 has been referred to as the “foie gras domain” after the zebrafish gene in which it was first analysed as it was noted to be particularly well conserved between species (Pfam domain PF11817) (Sadler et al., 2005). The N-terminal part of TRAPPC11 (residues 1-180) consists of four β-strands interspersed with four α-helices (Figure 4A). The C-terminal part (residues 567-1320) is less well resolved but forms the arm connecting to the vertex with TRAPPC12 and TRAPPC13 (Figure 2C).

**Figure 3.**
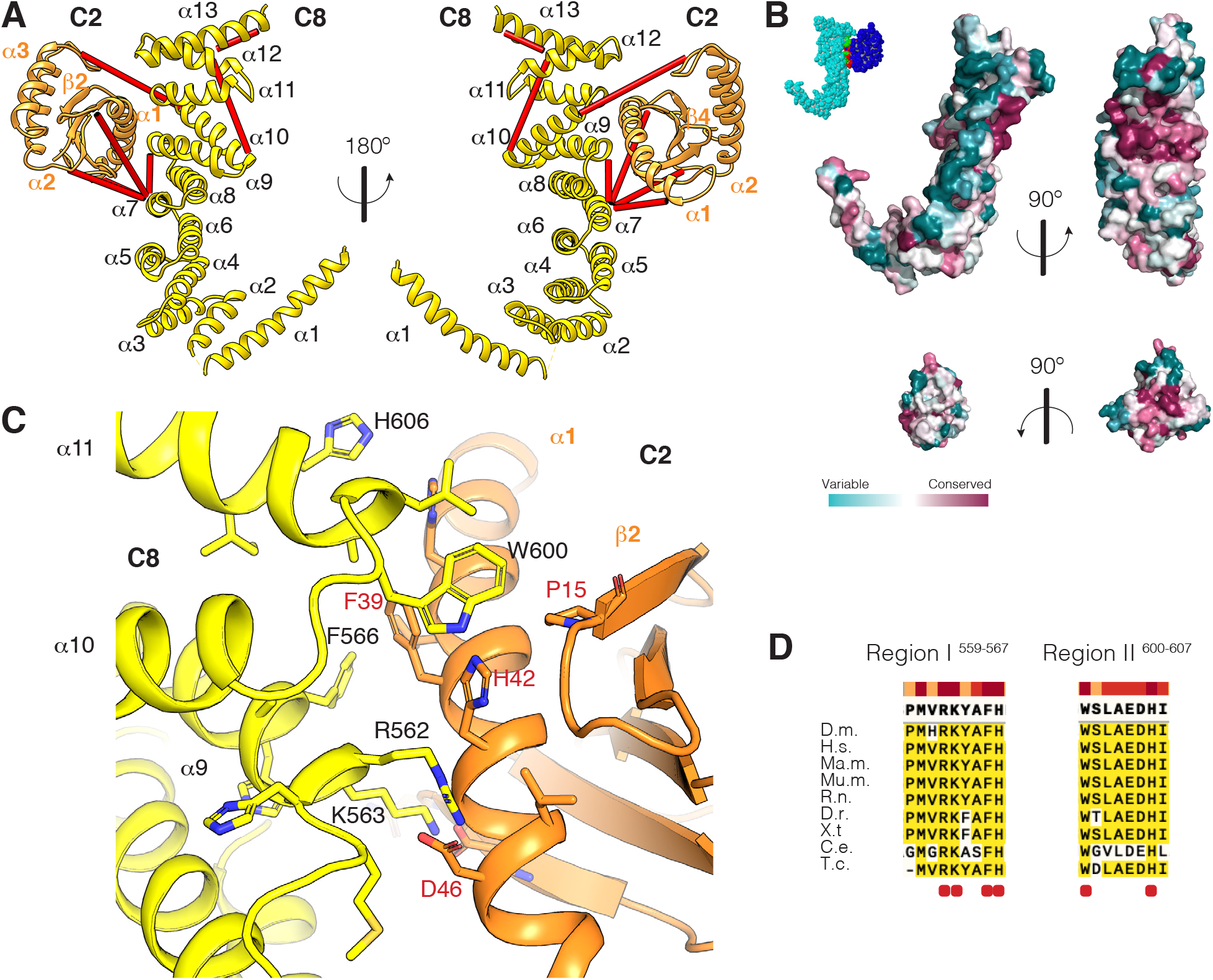
The TRAPPC2-TRAPPC8 interaction surface. (A) The TRAPPC2-TRAPPC8 subcomplex. TRAPPC8 (yellow) is formed by thirteen α helices and binds TRAPPC2 via helices 9, 10, 11. TRAPPC2 (orange) interacts with TRAPPC8 via α-helix 1 and the loop between β-strands 1 and 2. Cross-links mapped onto the model are shown as red lines. (B) Surface representation of TRAPPC8 and TRAPPC2 coloured according to evolutionary conservation as calculated using over orthologues identified with a Uniref 90 search with a 35% identity threshold and analysed by Consurf (Ashkenazy et al., 2016). Top: two views of TRAPPC8. The right one shows the surface of interaction between the two subunits. Bottom: two views of TRAPPC2. The surface of interaction is shown on the right. The inset at the top left corner shows a surface representation of the whole subcomplex for orientation purposes: TRAPPC8 is light blue, TRAPPC2 is dark blue, and the surface of interaction is coloured in red for the TRAPPC8 residues and green for the TRAPPC2 residues. (C) The TRAPPC2-TRAPPC8 interface. Structural model are coloured as in (A). Main residues involved in the interaction are shown as sticks. Labels for TRAPPC8 residues are black, for TRAPPC2 are red. (D) Alignment of the two TRAPPC8 conserved regions involved in the interaction with TRAPPC2. The residues highlighted in (C) are indicated with a red dot. Bar at the top indicates the degree of conservation.

**Figure 4.**
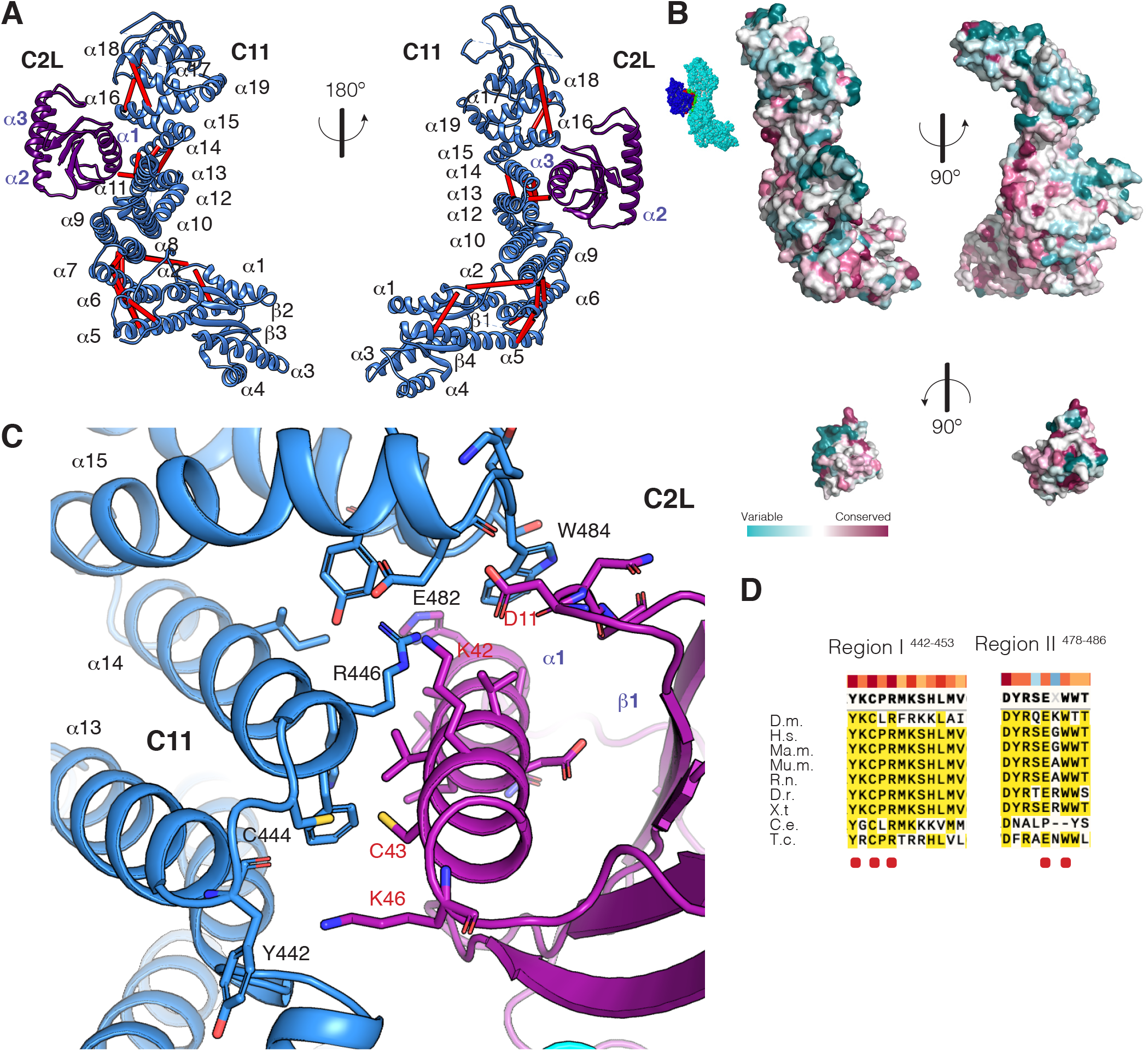
The TRAPPC2L-TRAPPC11 interaction surface. (A) Two views of the TRAPPC2L-TRAPPC11 subcomplex. The TRAPPC11 built model (blue) is formed by four β-strands and four α-helices at the N-terminal region that are joined to an α-solenoid comprising α-helices 5 to 18. α-helices 14 and 15 form the surface of interaction with TRAPPC2L (purple) and its surface of interaction involves α-helix 1 and the loop between β-strands 1 and 2. Cross-links mapped onto the model are shown as red lines. (B) Surface representation of TRAPPC11 and TRAPPC2L coloured according to evolutionary conservation (determined as for Figure 3B). Top: two views of TRAPPC11. The right one shows the surface of interaction between the two subunits. Bottom: two views of TRAPPC2L. The surface of interaction is shown on the right. The inset at the top left corner shows the whole subcomplex of TRAPPC11 (light blue) an TRAPPC2 (dark blue) with the surface of interaction coloured red for the TRAPPC11 residues and green for the TRAPPC2L residues. (C) The TRAPPC2L-TRAPPC11 interface. Structural model coloured as in (A). Main residues involved in the interaction are shown as sticks. Labels for TRAPPC11 residues are black, those for TRAPPC2L are red. (D) Alignment of the two TRAPPC8 conserved regions involved in the interaction with TRAPPC2. The residues highlighted in (C) are indicated with a red dot, the bar indicates the degree of conservation.

This proposed architecture of TRAPPC8 and TRAPPC11 binding to the core and also linking to a vertex occupied by TRAPPC12 and TRAPPC13 is further supported by the fact that these four TRAPPIII-specific subunits are able to form a stable subcomplex when co-expressed without the core subunits (Figure S6). Finally, the structural model of the core with the two flanking solenoids can be compared to the cross-linking data. We detected 146 total cross-links, and 75 of them mapped to residues present in the structural model - 32 cross-links were intermolecular and 43 intramolecular (Figure S7A). None of these cross-links exceeded the maximum distance constraint for disuccinimidyl dibutyric urea (DSBU) of ~30 Å, thus providing good validation for the atomic model of the TRAPPIII complex (Figure S7B and Table S1).

### TRAPPC2 and TRAPPC2L link the core to the arms of the complex in a similar manner

The primary interactions between the core and the arms are via binding of TRAPPC2 and TRAPPC2L to TRAPPC8 and TRAPPC11 respectively. The interaction between TRAPPC2 and TRAPPC8 encloses a total surface area of ~550 Å with the TRAPPC2 binding region of TRAPPC8 formed by α-helices 9 and 11 (Figure 3A). Conserved regions in TRAPPC8 (Pro559-His567 in α-helix 9 and Trp600-Ile607 in α-helix 11) form a hydrophobic pocket needed for the interaction (Figures 3B and 3C). In addition, α-helix 9 contains several polar and charged residues, such as Arg562 and Lys563, that are **l**ikely to interact with key residues in TRAPPC2, including Asp46 (Figures 3C and 3D). Interestingly, Asp46 appears to be particularly critical for the assembly of TRAPP complexes as mutation of this residue causes spondyloepiphyseal dysplasia tarda (SEDT) in humans (Gedeon et al., 1999; Sacher et al., 2019), and disrupts the TRAPP complexes in yeast (Brunet et al., 2013; Taussig et al., 2014; Zong et al., 2011). The second conserved region is located at the beginning of α-helix 11. It contributes to binding through interaction with residues in TRAPPC2 in α-helix 1 and to a lesser extent with residues located in the loop between β-strands 1and 2 (Figures 3C and 3D).

The surface of interaction between TRAPPC2L and TRAPPC11 is similar to that of TRAPPC2 and TRAPPC8, and has a total area of 558 Å (Figures 4A and 4B). The C-terminal α-helix of TRAPPC2L is shorter than that of TRAPPC2, and it interacts with the α-helices 14 and 15 of the TRAPPC11 α-solenoid (Figure 4C). We could identify two conserved regions in TRAPPC11 that are involved in the interaction. α-helix 14 (Tyr442 to Ile453), contacts TRAPPC2L between Asn33 and Lys42 (Figures 4C and 4D). This region is also conserved in TRAPPC2. The other region is between residues Asp478 to Thr486 at the end of α-helix 15, where residues such as Trp484 contact TRAPPC2L α helix 1 and the loop between β-strands 1 and 2, similar to TRAPPC8 and TRAPPC2 (Figures 4C and 4D).

### TRAPPC3 forms a second surface of interaction with TRAPPC8 and TRAPPC11

Two additional points of interaction between the arms and the core are visible in the TRAPPIII map as indicated by continuous densities between the flat surface of the core and the middle regions of both TRAPPC8 and TRAPPC11 (Figures S7C and S7D). TRAPPC8 is linked to density formed by the connection of the first two α-helices of TRAPPC3b and TRAPPC5. Similarly, density from TRAPPC11 connects to a region formed by the first two α-helices of TRAPPC3a and TRAPPC6. We could not build a model for TRAPPC8 or TRAPPC11 in these regions, but the cross-linking mass-spectrometry included cross-links between TRAPPC3a Lys41 and TRAPPC11 Lys649, and between the same lysine in TRAPPC3b and TRAPPC8 Ser1259 and Thr1263 (Figure 1E and Table S1).

### Location of the Rab1 binding site in the TRAPPIII structure

TRAPPIII activates Rab1 by catalysing the exchange of GDP for GTP, and then releasing the GTP-bound Rab1 to recruit effectors to the early secretory pathway. This exchange reaction is mediated by the central subunits of the core, and a crystal structure has been obtained for these subunits from yeast in a complex with a nucleotide-free form of Ypt1, the yeast orthologue of Rab1 (Cai et al., 2008). Rab1 is highly conserved through evolution, and all of the 22 residues of Ypt1 that were found to be within 4 Å of the interface with the yeast core are identical in *Drosophila* Rab1. Likewise, the residues of the core that bind Ypt1 correspond to the most highly conserved part of the surface of the *Drosophila* core (Figure 5A). We could thus model *Drosophila* Rab1 onto the equivalent region of the *Drosophila* core, and found that the GTPase fits into a space in the TRAPPIII density map (Figure 5B). Strikingly, in this position the surface of Rab1 is precisely abutted to the arm of TRAPPC8 that arches over the core before turning away to connect to the TRAPPC12/TRAPPC13 vertex. The part of Rab1 that contacts TRAPPC8 comprises two α-helixes, α3 and α4, of the canonical Rab structure (Pylypenko et al., 2018) (Figure 5B). Interestingly, one of these helixes contains one of the three Rab subfamily-specific sequences (RabSF3) that were defined as being conserved between Rabs of the same family but divergent between families (Moore et al., 1995; Pereira-Leal and Seabra, 2000). Thus, even though RabSF3 is located away from the switch regions that mediate binding to Rab1-specific effectors, its sequence is none-the-less specific to the Rab1 family. Therefore, the contact with TRAPPC8 could stabilise the interaction between Rab1 and the core and also increase specificity.

**Figure 5.**
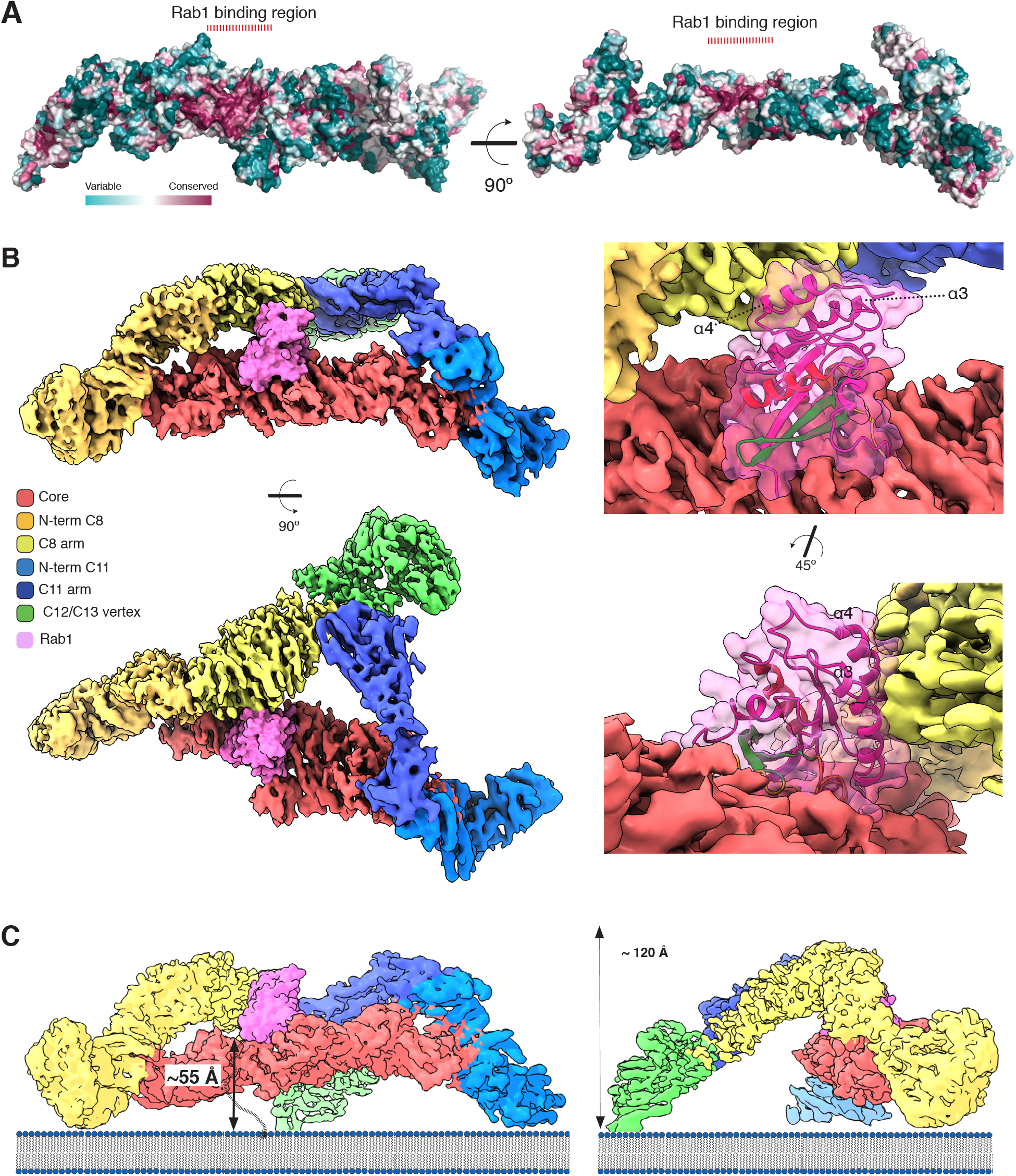
Rab1 binding site in TRAPPIII. (A) Left: Whole TRAPPIII model surface coloured by evolutionary conservation (determined as for Figure 3B). There is a highly conserved region in one of the flat surfaces of the core, which corresponds to the binding region of Ypt1 in complex with the central subunits of the yeast TRAPP core. (Cai et al., 2008). (B) Density map of Drosophila TRAPPIII with Rab1 (pink) modelled based on the location of Ypt1 bound to the core of yeast TRAPP. When bound to the core, Rab1 abuts the arm of TRAPPC8 (yellow). Enlargements with the Rab1 ribbon structures showing canonical Rab helices α3 and α4 are facing the surface of TRAPPC8. (C) Model of TRAPPIII with bound Rab1 on a flat lipid membrane. The distance between the of the Rab1-binding site and the membrane surface is 55 Å, shorter that the predicted maximum length of the unstructured hypervariable domain of Rab1 that connects it to its C-terminal lipid anchor (shown as hashed line), and on a curved bilayer such as a vesicle the distance would be shorter.

GDP-bound forms of Rabs bind the cytosolic chaperone GDP-dissociation inhibitor (GDI) that masks the C-terminal prenyl groups. This enables the Rab to be soluble in the cytosol, and activation of Rabs is believed to occur after GDI has released the GDP-bound form on to a membrane. This means that GEFs like TRAPP act on the membrane rather than in the cytosol (Barr, 2013; Blümer et al., 2013; Pylypenko et al., 2006). Consistent with this, the TRAPPs have greater activity toward Rabs bound to liposomes, which indicates that the TRAPPs interact with the membrane surface, an interaction that could be promoted in vivo by other proteins present on the membrane (Riedel et al., 2017; Thomas and Fromme, 2016). Therefore, in addition to allowing Rab1 to access the catalytic site in the core, the structure of TRAPP needs to be compatible with the substrate Rab1 being connected to a lipid bilayer via the unstructured hypervariable domain that links the GTPase to the prenyl groups that mediate membrane attachment (Li et al., 2014). The location of the hypervariable domain when Rab1 is bound to TRAPP is unknown as it was not included in the form of Rab1 used to generate a crystal structure with the core subunits. However, modelling the TRAPPIII structure on a flat surface places the Rab1 binding site 55 Å above this surface (Figure 5C), a distance that would be readily accommodated by the ~95 Å that the 27 residue hypervariable domain of *Drosophila* Rab1 (Gly177-Gly203) could reach at its maximum extent. At present, it is formally possible that the complex is positioned perpendicular to the membrane, which would move the Rab binding site closer, but we can at least say that earlier proposals that the sides or the underneath of the core could contact the membrane are not sterically possible as the vertices extend beyond the core (Cai et al., 2008; Kim et al., 2006), whereas, having all three vertices on the membrane would be compatible with exchange activity.

Finally, we investigated the relevance of the flexibility of TRAPPIII for Rab1 binding. As noted above, the limits on the resolution of the density map implied that the complex is not entirely rigid. The best resolved part of the map is the core and the associated regions of TRAPPC8 and TRAPPC11. To assess its movement relative to the rest of the complex we used multibody refinement to look at variation of the particles within the dataset and analysed the results by principle component analysis (Nakane et al., 2018). Almost half of the variability between particles can be accounted for by a movement vector corresponding to a rocking of the arms relative to the core (Figures S8A and S8B, and Video S1). This indicates that the arms have sufficiently flexibility for TRAPPC8 to move over the Rab1 binding site to the point that it would block binding of the GTPase to the catalytic site on the core (Figure S8C). This provides a possible mechanism by which the interactions formed by the four subunits of the arms could regulate exchange activity.

### Architecture of the TRAPPII complex

The TRAPPII complex shares the core subunits with TRAPPIII but has different additional subunits which allow it to activate Rab11 (Figure 1A). To compare the overall architecture of the two complexes we expressed a recombinant form of Drosophila TRAPPII and subjected it to single particle cryo-EM imaging. A low resolution 3D map shows that the overall architecture of TRAPPII is similar to that of TRAPPIII, with an elongated rod of the dimensions of the core attached to two arms that connect at their opposite ends to form an irregular triangle (Figure S9A). Application of cross-linking mass spectrometry indicates that TRAPPC9 is linked to the core through TRAPPC2, and TRAPPC10 is linked via TRAPPC2L, with the latter showing a cross link via the same Lys47 residue that linked to TRAPPC11 in TRAPPIII (Figures S9B and S9C, Table S1). This is consistent with studies in yeast where the TRAPPC2L orthologue Tca17 is required for association of Trs130 with the TRAPPII complex (Choi et al., 2011; Milev et al., 2018). The pattern of cross-linking between the core subunits is similar to that found in TRAPPIII, and TRAPPC3 Lys 41 and TRAPPC6 Lys 104 also links to TRAPPC10, analogous to the links these core subunits form to TRAPPC11. Together, these findings show that both metazoan TRAPP complexes share an architecture that consists of a central core held between two elongated arms.

## Discussion

The TRAPP complexes have emerged as arguably the two most critical activators of Golgi Rab function, with Rab1 acting as the master regulator of entry into the early compartments of the stack, as well as being a key player in autophagy. Biochemical and structural studies have elegantly shown that Rab activation in vitro requires only three of the small subunits at the core of these large structures (Cai et al., 2008; Kim et al., 2006). The presence of further, and larger, subunits presumably reflects the need for precise temporal and spatial control of the activation of these essential GTPases, but how these subunits might exert control over the core GEF activity has been unclear. The cryo-EM structure of the entire TRAPPIII complex presented here clearly shows how the arms of the complex are attached to the core and resolves long standing uncertainty on this issue. Previous structural and genetic studies have provided unambiguous evidence that TRAPPC8 binds to TRAPPC2 which is present at one end of the core (Brunet et al., 2013; Kim et al., 2006; Pinar et al., 2019; Tan et al., 2013). Likewise, it is clear that in TRAPPII, TRAPPC9 binds to the same subunit. However, the situation for the other arms has been less clear as TRAPPC2L is absent from yeast TRAPPIII, and there is no crystal structure of a complex between TRAPPC2L and other core subunits. It has been proposed that TRAPPC2L acts in TRAPPII in yeast to attach TRAPPC11 to the core, but this has not been universally accepted (Choi et al., 2011; Lipatova and Segev, 2019; Montpetit and Conibear, 2009). Our results unambiguously place TRAPPC2L at the opposite end of the core from TRAPPC2 and show how it attaches to TRAPPC11. TRAPPC2L is likely to have the equivalent position in TRAPPII so as to attach TRAPPC10. Interestingly, in *Drosophila* or fungal mutants lacking TRAPPC10, TRAPPC2L appears to dissociate from the core, suggesting its binding is stabilised by the interaction, possibly by virtue of the triangulation to the other end of the core via the other arm (Pinar et al., 2019; Riedel et al., 2017). Alternatively, quality control mechanisms in the cell may recognise the partly assembled complex and degrade some of the subunits. The fact that the related TRAPP2C and TRAPPC2L subunits have equivalent roles at opposite ends of the complex is echoed by the fact that the sequence of TRAPPC11 is distantly related to that of TRAPPC10, with the same being true for TRAPPC8 and TRAPPC9 that bind TRAPPC2 (Scrivens et al., 2011; Wendler et al., 2010). This indicates that a eukaryotic precursor had a single ur-TRAPP, and gene duplication gave rise to the two TRAPPs that appear to have been present in the last eukaryotic common ancestor.

The TRAPPIII structure also reveals the location of TRAPPC12 and TRAPPC13 in the complex, showing that they are present at the joint between the TRAPPC8 and TRAPPC11 arms. Unlike, the arms themselves, these subunits do not seem to have equivalents in *Drosophila* TRAPPII. The TRAPPII complex of budding yeast has an additional subunit, Trs65, that was originally proposed to be a yeast homolog of TRAPPC13, but now appears to be a relative that arose by duplication in fungi, with some budding yeast then losing TRAPPC13 itself, along with TRAPPC12 and TRAPPC11 (Choi et al., 2011; Pinar et al., 2019; Riedel et al., 2017). Interestingly, something similar seems to have happened in some other phyla with a TRAPPC13 relative being recently found to associate with at least a subset of TRAPPII in plants (TRIPP), and vertebrates (C7orf43/TRAPPC14), suggesting that the vertex of the TRAPP arms is a convenient place to bolt on additional subunits (Cuenca et al., 2019; Garcia et al., 2020).

Irrespective of structural and evolutionary relationships, the key question that remains is the role of these large arms on the TRAPP complex. The structure of TRAPPIII reveals how the arms could serve as regulators despite being attached via subunits that are distal to those that mediate GEF activity. Like other GEFs, TRAPPIII is likely to be regulated by recruitment to the membranes where it can access GDP-bound Rab1 (Barr, 2013; Blümer et al., 2013). The size of the arms provides a large surface area that components of membrane traffic or autophagy could bind to without sterically inhibiting exchange activity, and indeed interactions have been reported between TRAPPs and a wide range of potential regulators including the Sec23/Sec24 and the Sec13/31 subunits of the COPII coat, the COPI coat, Arf1 exchange factors, the Rab GAP TBC1D14 and the autophagy proteins Atg2 and Atg9 (Kakuta et al., 2012; Lamb et al., 2016; Stanga et al., 2019; Tan et al., 2013). In addition, yeast TRAPPII can be activated in vitro by the small GTPase Arf1 (Thomas et al., 2018). However, the architecture of the TRAPPIII complex indicates that the arms could also have more direct effects on activation. Firstly, TRAPPC8 is positioned such that it would contact Rab1 bound to the GEF active site on the core, and this could both enhance the rate of exchange and also improve selectivity for Rab1 over other Rabs. Secondly, the flexibility of the arms is such that TRAPPC8 could move so as to interfere with, rather than augment, access to the active site, and therefore interactions that moved the arms could alter the activity of membrane-bound TRAPPIII. Finally, we observe apparent contacts between both arms and the TRAPPC3 subunits near the centre of the core which raiases the possibility of allosteric regulation. Clearly further work will be required to address the in vivo significance of these various possible modes of regulation, but hopefully the architecture reported here will greatly facilitate this by guiding the construction of specific alterations to the complex.

## Methods

### Expression and purification of TRAPP Complexes

The TRAPP purification protocol is based on previous work (Riedel et al., 2017). The complexes were expressed in insect cells (Sf9 or Hi5 lines) using the MultiBac System (Nie et al., 2014). A pACEBac1 plasmid containing the seven *Drosophila melanogaster* core subunits, pACEBac1-C1-C6, was fused using Cre recombinase (New England Biolabs), to a pIDS vector containing TRAPPC9 and TRAPPC10 to generate the plasmid pACEBac1-TRAPPII-complete. A similar strategy was followed to construct the pACEBac1-TRAPPIII-complete: TRAPPC8 and TRAPPC11 were cloned into pIDS, and TRAPPC12 and TRAPPC13 were cloned into pIDC. The resulting plasmids were recombined into pACEBac1-C1-C6 vector to express miniTRAPPIII, with only TRAPPC8 and TRAPPC11, or the complete TRAPPIII (Riedel et al., 2017). The TRAPPC3 subunit was tagged with Strep-TagII, and TRAPPC11 (TRAPPIII) or TRAPPC10 (TRAPPII) were FLAG-tagged, both at the N-terminus. An additional pACEBac1-C1-C6 with the TRAPPC2L subunit tagged with the ZZ domain at the N-terminus was used for the expression of the TRAPP core. The linker sequence between each subunit and the tag included a site for the HRV-3C protease. The pIDS and pIDC plasmids containing the specific TRAPP subunits were also fused to an empty pACEBac1 to express these subunits in the absence of the TRAPP core subunits.

Bacmids were made using the EMBacY system (Nie et al., 2014). A 500 ml suspension of Sf9 cells (2×10^6^ cells/ml) was infected with 5 ml of fresh P2 baculovirus and incubated at 27°C and 124 rpm. Cells co-expressing the TRAPP core, TRAPPII or TRAPPIII were harvested after 66 hours (at 75-80% viability) by centrifugation at 2250 × g for 10min at 4°C. Pellet was washed once with PBS, centrifuged again, and processed immediately for the whole TRAPP complexes, or kept at −80°C in the case of the TRAPP core. Initially, pellets were resuspended in Buffer A (50 mM Hepes-KOH pH 7.44, 150 mM KAc, 1 mM DTT, 0.1% IGEPAL CA-630) with inhibitors (1mM PMFS, cOmpleteTM, 0.4 μM pepstatin, 0.24 μM leupeptin, 5 μM MG132) at a ratio of 30 ml per 500 ml of initial culture. The cell suspension was vortexed and incubated at 4°C for 10 min, before lysis by 15-20 strokes of a tight-fitting dounce homogeniser. The lysate was clarified by centrifugation at 32000 × g for 30 min at 4°C. Cleared lysate was mixed with the appropriate equilibrated slurry: Strep-Tactin Superflow plus (Quiagen) (400 μl per 500 ml of initial culture), Anti-Flag M2 affinity gel (Sigma) (100 μl per 500 ml culture) or IgG Sepharose (6 Fast Flow, GE Healthcare) (500 μl per 500 ml culture), and incubated on rotation wheel for one hour at 4°C. Beads were washed three times with ten bead volumes of Buffer A plus 0.05% IGEPAL CA-630. Bound material was eluted by washing the beads with five bead volumes of Buffer A containing either 100 μg/ml FLAG peptide (anti-FLAG) or 2.5 mM desthiobiotin (Strep-Tactin). The eluted fraction was analysed by SDS-PAGE, concentrated and buffer exchanged. Alternatively, the bound complexes were eluted by tag cleavage incubating the slurry with PreScission protease (~ 10U/ml) overnight at 4°C. The eluted solution was mixed with glutathione-Sepharose to remove the PreScission protease.

The TRAPP core complex was purified further by gel filtration (SEC) using Superose 10/30 (GE Healthcare) (Figure S6) for small samples, or Superdex200 16/100 equilibrated in Buffer A plus 0.005% IGEPAL CA-630. This protocol was escalated to 6 litre cultures (12 × 500 ml) for TRAPPII, TRAPPIII and miniTRAPPIII. We found that increasing the KAc concentration prevents the formation of aggregates during SEC step purification and so the composition of Buffer A composition was adjusted to 250 mM KAc, and the IGEPAL CA-630 removed during the subsequent bead washing. Detergent-free samples were concentrated up to 3-5 mg/ml, and ~100μl fractions were loaded onto a TSKgel G4000SWXL column (TOHO Bioscience) in 50mM Hepes-KOH pH 7.44, 250 mM KAc, 1mM DTT (Buffer B). Eluted peaks were collected in 100 μl fractions and analyzed by SDS-PAGE and Coomassie staining (Figure S1A). Protein identification by mass spectrometry was used to assess the integrity of the purified complexes. The best yield for the purification of whole TRAPP complexes was obtained using the TRAPPC10 or TRAPPC11 Flag-tagged subunits as baits for the affinity chromatography. There was no difference in stoichiometry or in vitro GEF activity between complexes obtained by FLAG peptide elution or by HRV-3C protease cleavage, and so we continued with the former method.

### SEC-MALs and iSCAT analysis

For SEC-MALs, purified TRAPPIII and miniTRAPPIII (~100 μl at 0.5 mg/ml) were resolved on a Superose 6 10/300 column (GE Healthcare) in Buffer B, with a flow rate of 0.5 ml/min. Protein was detected with 280 nm UV light (Agilent Technology 1260), a quasielastic light scattering module (DAWN-8+, Wyatt Technology), and a differential refractometer (Optilab T-rEX, Wyatt Technology). Molar masses of peaks in the elution profile were calculated from the light scattering and protein concentration, quantified using the differential refractive index of the peak, assuming dn/dc = 0.186, with ASTRA7 (Wyatt Technology).

For iSCAT, TRAPPIII, TRAPPII and miniTRAPPIII were diluted to a final concentration of 25, 50 and 100 nM irrespectively in Buffer B. 10μl of each sample was applied to 10μl of Buffer B on a cleaned glass coverslip (n° 1.5, 24 × 50 mm) coated with a silicone well frame and analysed for 10 min at a rate of 600 frames/ min with an ONEMP mass photometer (Refeyn LTD, Oxford, UK). 25 nM and 50 nM BSA solution were used as standards for calibration. For each recording of a BSA standard, a histogram was made and fitted with Gaussians according to how many peaks are resolved. Fitted centres of these Gaussians and the corresponding masses that they are assigned to were plotted and fitted to a straight line. The resulting parameters were used as conversion between measured contrast and mass for the TRAPP samples (Cole et al., 2017). Data were acquired and analyzed using AcquireMP and DiscoverMP (v1.2.3) (Refeyn LTD, v1.1.3). Measurements were repeated at 4°C and RT with similar results.

### Negative-stain EM

After gel filtration, TRAPPIII samples were diluted to 0.008-0.009 mg/ml (~16-20 μM) and applied to EM grids. 3 μl of diluted sample was deposited onto a glow-discharged (Edwards S150B, 30sec, 30-50 mA, 1,2kV, 10-2 mbar) continuous carbon grid (CF400-CU-UL, Electron Microscopy Sciences). After one minute at RT, the grid was blotted and washed by immersion in a 100 μl drop of fresh 2% uranyl acetate and blotted again. Then, the grid was stained by two rounds of 2% uranyl acetate immersion for 30 sec and blotting, before being air-dried. Micrographs were collected on a Tecnai T12 microscope (Thermo Fisher Scientific) operating at 120 keV with a tungsten electron source and a 2k × 2k CCD camera (Orius SC200W, Gatan, Inc.). Nominal magnification was 15,000×, giving a 3.50 Å/pixel sampling at the object level. Images were collected with a dose of 50 e−/Å2, and a nominal defocus of −1 μm. In total, 100 micrographs were collected. TRAPPIII particles were manually picked and subjected to initial 2D classification using Relion v3.0 (Zivanov et al., 2018). Automated particle picking was made using EMAN2, and particles coordinates were imported into Relion v3.0. The initial 22,986 particles were subjected to two rounds of Class2D classification resulting in 25 class averages. 3123 particles were sorted to build a 3D initial model de novo. This model was used as a reference map for 3D refinement of the total subset of 10,077 good quality particles. The final model was obtained after Class3D classification and another round of 3D refinement.

### Cryo-EM grid preparation

After gel filtration, TRAPPIII, TRAPPII or miniTRAPPIII were diluted to 0.9-1 mM (0.5 mg/ml) in buffer supplemented with IGEPAL CA-630 to reach a final concentration of 0.005%. Samples were applied to freshly glow-discharged (Edwards S150B, 45 sec, 30-50 mA, 1,2kV, 10-2 mbar) copper holey carbon grids (Quantifoil, Cu-R1.2/1.3) under 100% humidity. Excess sample was blotted away and the grids were subsequently plunge-frozen in liquid ethane using a Vitrobot Mark III (Thermo Fisher Scientific).

### Data collection

#### TRAPPIII

A total of 3671 movies were recorded on a Titan Krios electron microscope (Thermo Fisher Scientific – FEI) operating at 300 kW with a calibrated magnification of 75000x and corresponding to a magnified pixel size of 1.04 Å. Micrographs were recorded using a Falcon III direct electron detector in counting mode with a dose rate of ~ 0.5 e/Å2/s and defocus ranging from −2.2 μm to −4 μm. The total exposure time was 60 s, and intermediate frames were recorded in 0.8 s intervals, resulting in an accumulated dose of ~30 e/Å2 and a total of 75 frames per micrographs.1190 movies out of the 3671 datasets were collected with the stage titled at 19°, this angle chosen according to the output from the cryoEF algorithm (Naydenova and Russo, 2017).

#### MiniTRAPPIII

A small dataset of 385 micrographs was acquired under the same conditions described for TRAPPIII. Data derived from these micrographs were used for building a partial ab initio 3D model used as a reference map for TRAPPIII and miniTRAPPIII.

A second dataset of 3,443 micrographs was acquired on a Titan Krios EM operating at 300 kW with a calibrated magnification of 105000x and corresponding to a magnified pixel size of 1.047 Å. Micrographs were recorded using a K2 direct electron detector (Gatan) equipped with a Cs corrector and an energy filter. Images were collected over 12 s in counting mode with 0.3 s (~e/Å2/s) frame time and a slit width of 20 eV. The total exposure was 45.6 e/Å2, and the defocus ranged from −1.5 μm to −2.5 μm.

#### TRAPPII

A total of 364 micrographs were recorded on a Titan Krios III EM operating at 300 kW with a calibrated magnification of 75000x and corresponding to a magnified pixel size of 1.09 Å. Settings for the acquisition were similar to those for TRAPPIII (Falcon III in counting mode, ~ 0.5 e/Å2/s, defocus −2.2 μm to −4 μm, exposure 60s, total dose~30 e/Å2).

### Image processing

Dose fractionated image stacks were subjected to beam-induced motion correction and filtered according to the exposure dose using MotionCor2 (Zheng et al., 2017). The sum of each movie was applied to CTF parameters determination by CTFFIND 4.1 (Rohou and Grigorieff, 2015). For the tilted data, the CTF was corrected according to the focus gradient of each particle using goCTF (Su, 2019). A customize script was written to run the goCTF v1.2.0 software in batch. Particles were picked using cryOLO 1.5 (Wagner et al., 2019). Particles from the small miniTRAPPIII dataset were subjected to 2D classification, and 22,897 particles were chosen to create an ab initio 3D model using the Frealign tool implemented in cisTEM (Grant et al., 2018; Grigorieff, 2016). This resulted in a rough 3D map at 7 Å used for 3D classification of particles from the larger TRAPPIII and the miniTRAPPIII datasets. In the case of TRAPPIII, tilted and non-tilted data were CTF corrected, extracted and subjected to two rounds of reference-free 2D classification using Relion 3.1. Selected 2D classes were used for a 3D classification resulting in three different classes that were similar among the different datasets. The corresponding particles to the cleanest 3D class from each dataset were joined, reextracted and subjected to an additional round of 2D classification. The selected particles after this round were 3D refined. After this, 353,400 particles were subjected to 3D masked refinement followed by map sharpening in Relion 3. The estimated CTF parameters were refined, and per-particle reference-based beam-induced motion correction was performed using Bayesian polishing. The final map has a global resolution of 5.8 Å. Reported resolution is based on the gold-standard Fourier shell correlation (FSC) using the 0.143 criteria. Local resolution was estimated using the Relion 3.1 implementation. A similar strategy was followed for miniTRAPPIII, but the global resolution was higher than the consensus map for TRAPPIII, at 4 Å, but with a higher range in the local resolution. This is due to the higher number of micrographs and the strong preferential orientation of this complex (EOD 0.62 (Naydenova and Russo, 2017)) vIn the case of TRAPPII, 43,161 particles were picked using crYOLO. After several runs of 2D classification, 22,570 particles were selected to generate a rough ab initio 3D model. This model was used as a reference map for a 3D classification. Particles corresponding to the best 3D classes were joined and subjected to 2D classification, and 3084 good particles were selected to generate a 3D model at 15 Å resolution.

### Multi-body refinement

To improve the density, increase the resolution and characterize the conformational dynamics, we performed multi-body refinement with RELION 3.1 (Nakane et al., 2018). TRAPPIII was divided into three or four discrete bodies composed initially by the whole flat rod, the TRAPPC11 arms, and the TRAPPC8 arms plus the TRAPPC12-TRAPPC13 vertex, with the latter being isolated as an additional body for the four bodies approach. In later trials, the core alone constituted one body, and the whole TRAPPC11 and the whole TRAPPC8 plus TRAPPC12 and TRAPPC13 the other two. Masks for multi-body refinement were made in UCSF Chimera 1.15 from the consensus map (Pettersen et al., 2004). The standard deviation of the Gaussian prior on the rotations was set to 10 degrees for all three bodies. The standard deviations on the body translations were all set to two pixels. The maps for the three discrete bodies after multi-body refinement were post-processed individually and combined using Phenix (Liebschner et al., 2019). There was an increase in resolution (Figure S4, body1: core 4.2 Å, body2: C8-C12-C13 4.4 Å, and body3: C11 5.5 Å), enabling interpretation of the density for the N-terminal regions of TRAPPC8 and TRAPPC11.

### Flexibility analysis

We used the relion_flex_analyse program to perform a principal component analysis (PCA) on the relative orientations of the bodies of a subset of 110,367 particles (Zivanov et al., 2018). The PCA is performed on six variables per body (3 translations and 3 rotations). We analysed the variance in the rotations and translation of the bodies explained by the different eigenvectors. UCSF Chimera 1.15 was used to generate movies of the reconstructed body densities repositioned along these eigenvectors. Individual maps of the bodies, positioned relative to each other according to the rotations and translations corresponding to the centre of the amplitude along the different eigenvectors, were used to calculate the rotation angles and the translation distances (Pettersen et al., 2004).

### Model building and refinement

The *Drosophila* TRAPP core subunits were modelled using Modeller (Sali and Blundell, 1993; Webb and Sali, 2016). Previously reported crystal structures for the subunits were used as homology models. (1HQ3 (Jang et al., 2002); 2J3T and 2J3W (Kim et al., 2006), 3PR6 (Wang et al., 2014)). The core subunit models were initially fitted into the maps using UCSF Chimera 1.15 and the chains were manually adjusted in Coot 0.9 (Burnley et al., 2017; Casañal et al., 2020; Pettersen et al., 2004). The final models were then refined in Phenix within the real-space refinement module, using secondary structure and Ramachandran restraints (Liebschner et al., 2019). The TRAPPC8 and TRAPPC11 N-terminal regions were built de novo. Initial models, generated using trRosetta (Yang et al., 2020), were docked into the corresponding map and manually adjusted in Coot 0.9 (Casañal et al., 2020). Regions in which the sequence could be unambiguously docked and/or supported by cross-linking data were built and kept in the final models, which were refined against the whole maps and evaluated in Phenix (Liebschner et al., 2019). Figures were generated using PyMOL (Version 2.0 Schrödinger, LLC), UCSF Chimera 1.15 and UCSF Chimera X (Pettersen et al., 2004; 2020). Model geometry evaluation and half-map validation were performed using Molprobity (Williams et al., 2018). The final refinement statistics are provided in Table 1.

### Cross-linking coupled to mass spectrometry (XL-MS)

300 μl of TRAPPIII and TRAPPII in Buffer B at ~ 0.8-1 mg/ml (1.8 ~ 2 mM) were cross-linked with the N-hydroxysuccinimide (NHS) ester disuccinimidyl dibutyric urea (DSBU, formerly BuUrBu). Cross-linking was at 45 min at room temperature at 150 times the protein concentration, and then quenched by the addition of NH_4_HCO_3_ to a final concentration of 50 mM, and incubating for 15 min. The cross-linked samples were precipitated with methanol/chloroform (Wessel and Flügge, 1984), resuspended in 8M urea, reduced with 10 mM DTT and alkylated with 50 mM iodoacetamide. Following alkylation, proteins were diluted with 50 mM NH4HCO3 to a final concentration of 2M urea and digested with trypsin (Promega, UK), at an enzyme-to-substrate ratio of 1:20, overnight at 37°C or sequentially with trypsin and Glu-C (Promega, UK) at an enzyme-to-substrate ratio of 1:20 and 1:50 at 37°C and 25°C, respectively. The samples were acidified with formic acid to a final concentration of 2% (v/v), then split into two equal amounts for peptide fractionation by peptide size exclusion and reverse phase C18 high pH chromatography (C18-Hi-pH). For peptide size exclusion, a Superdex Peptide 3.2/300 column (GE Healthcare) with 30% (v/v) acetonitrile/0.1% (v/v) TFA as mobile phase and a flow rate of 50 μl min−1 was used, and fractions collected every 2 minutes over the elution volume of 1.0 ml to 1.7 ml. C18-Hi-pH fractionation was carried out on an Acquity UPLC CSH C18 1.7 μm, 1.0 × 100 mm column (Waters) over a gradient of acetonitrile 2-40% (v/v) and ammonium hydrogen bicarbonate 100 mM.

The fractions were lyophilized and resuspended in 2% (v/v) acetonitrile and 2% (v/v) formic acid and analyzed by nano-scale capillary LC-MS/MS using an Ultimate U3000 HPLC (ThermoScientific Dionex, USA) to deliver a flow of approximately 300 nl.min−1. A C18 Acclaim PepMap100 5 μm, 100 μm × 20 mm nanoViper (ThermoScientific Dionex, USA), trapped the peptides before separation on a C18 Acclaim PepMap100 3 μm, 75 μm × 250 mm nanoViper (ThermoScientific Dionex, USA). Peptides were eluted with a gradient of acetonitrile. The analytical column outlet was directly interfaced via a nano-flow electrospray ionization source, with a hybrid quadrupole orbitrap mass spectrometer (Orbitrap Q-Exactive HF-X, Thermo Scientific). MS data were acquired in data-dependent mode. High-resolution full scans (R=120,000, m/z 350-2000) were recorded in the Orbitrap followed by higher energy collision dissociation (HCD, stepped collision energy 30 ± 3) of the 10 most intense MS peaks. MS/MS scans (R=45,000) were acquired with a dynamic exclusion window of 20s being applied.

For data analysis, Xcalibur raw files were converted into the MGF format by MSConvert (Proteowizard) and put into MeroX (Götze et al., 2012; Kessner et al., 2008). Searches were performed against an ad hoc protein database containing the sequences of the complexes and a set of randomized decoy sequences generated by the software. The following parameters were set for the searches: a maximum number of missed cleavages of three; targeted residues K, S, Y and T; minimum peptide length of five amino acids; variable modifications: carbamidomethyl-Cys (mass shift 57.02146 Da), Met-oxidation (mass shift 15.99491 Da); DSBU modification fragments: 85.05276 Da and 111.03203 (precision: 5 ppm MS1 and 10 ppm MS2); false discovery rate cut-off: 5%. Finally, each fragmentation spectra was manually inspected and validated. Data were analyzed and figures generated using xiView (github.com/Rappsilber-Laboratory/xiView) and Xlink Analyzer (Kosinski et al., 2015).

### GEF activity assays

GEF assays were performed as previously (Riedel et al., 2017). In summary, the activity on His-tagged Rabs was determined by the exchange of nonfluorescent GTP for mant-GDP using a PHERASTAR plate reader. All Rabs, and TRAPP complexes were buffer exchanged into HKM (20 mM Hepes-KOH, pH 7.4, 150 mM KOAc, 2 mM MgCl2, and 1 mM DTT). Reactions containing 250 nM mant-GDP–labelled Rab alone, Rab and 200 *μ*M GTP, or adding to the mix 10 mM EDTA, or 25 nM of the corresponding GEF, were set up in 96-well black-bottomed plates (Corning), and fluorescence decay was measured at 30°C.

## Acknowledgements

We thank Giuseppe Cannone, Grigory Sharov and Anna Yeates from the MRC LMB, and the eBIC Diamond staff, for assistance in cryo-EM data collection; Stephen McLaughlin and Chris Johnson for help with biophysics, and Jake Grimmett and Toby Darling for computational support. We are indebted to Ester Vazquéz and Ana Casañal for advice on handling cryo-EM samples and helping in data collection, Pavel Afanasyev and Arka Chakraborty for advice on image processing. We are grateful to Andrea Nans, Tim Stevens, Elyse Fischer, Alba Herrero and Ana Torroja for help with goCTF and CryoSParc software, and to Alison Gillingham and Jérôme Cattin-Ortolá for comments on the text. This study made use of electron microscopes at the MRC LMB EM Facility and the UK’s national Electron Bio-imaging Centre (eBIC) under proposal EM17434 funded by MRC. All other funding was from the MRC (File reference number MC_U105178783).

## Data and materials availability

The final reconstructed maps from each frame and the weighted sum are deposited in the Electron Microscopy Data Bank (TRAPPIII consensus map: EMD-12056, miniTRAPPIII consensus map: EMD-12063, body1-Core: EMD-12052, body2-C8/C12/C13: EMD-12053, body3-C11: EMD-12054, TRAPPII: EMD12066), and the refined atomic model in the Protein Data Bank (TRAPPIII: 7B6R, MiniTRAPPIII: 7B7O, TRAPP Core: 7B6D, C8: 7B6E, C11: 7B6H).

## Author contributions

A.G. and S.M. devised the study, A.G. performed the biochemistry, the single particle cryo-EM, and analysed the cross-linking mass spectrometry. V.J.P.-H. assisted with model building, G.D. performed and analysed the cross-linking mass spectrometry, and A.G. and S.M. wrote the manuscript.

## Supplemental Figures

**Figure S1.**
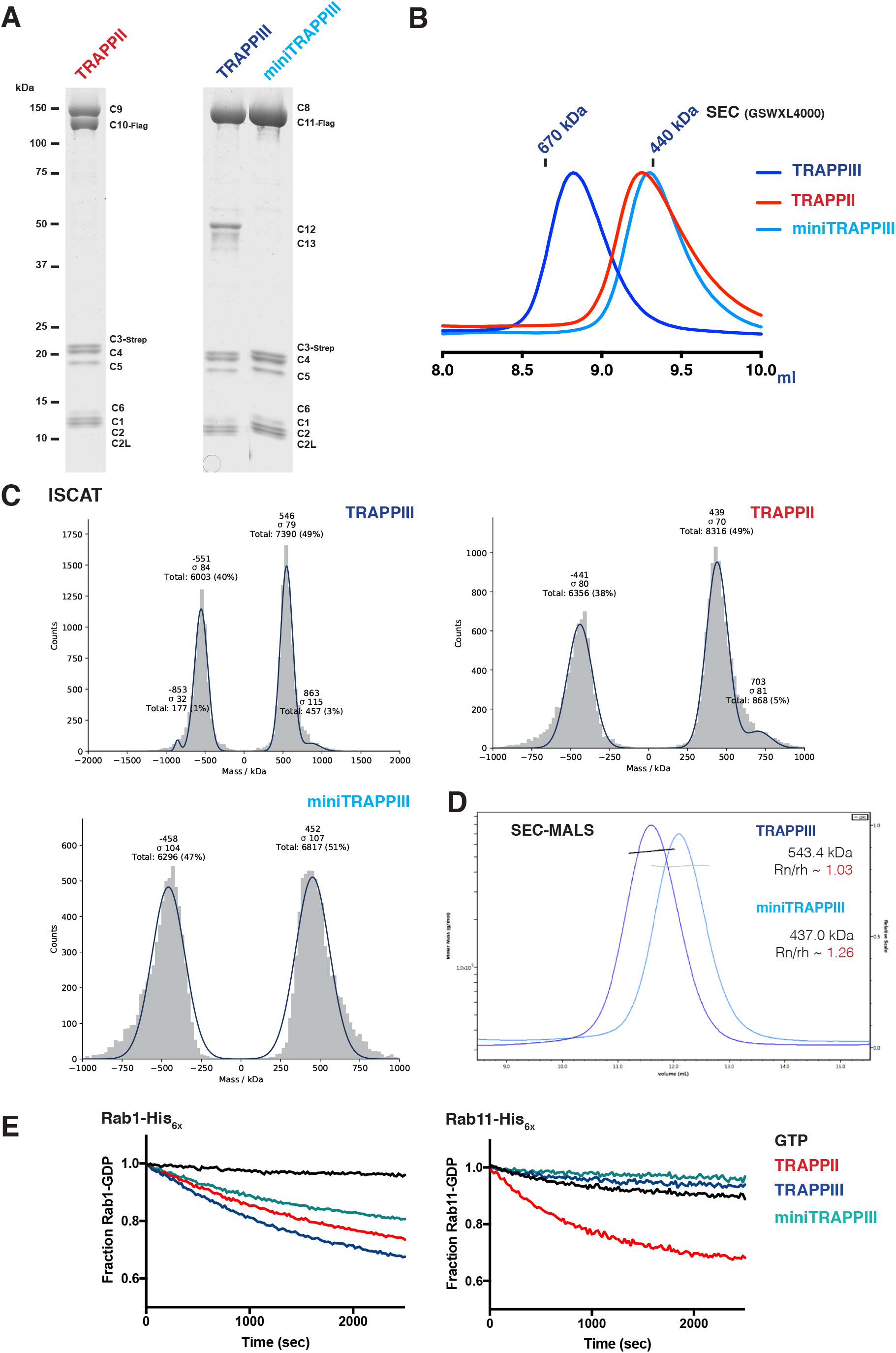
Characterization of the TRAPP complexes. (A) Coomassie blue–stained protein gels of recombinant *Drosophila* TRAPP complexes. The different TRAPP subunits are indicated. (B) Normalized UV traces from SEC of the TRAPP complexes. (C) iSCAT analysis of 50 nM samples of TRAPPIII, TRAPPII and miniTRAPPIII. Movies for each sample were recorded and their contrast translated into histograms. The histograms were fitted to a Gaussian distribution and compared with measurements from BSA standards to convert contrast into molecular mass (Cole et al., 2017) (TRAPPIII ≈ 548.5 kDa; TRAPPII ≈ 440 kDa; miniTRAPPIII ≈455 kDa). (D) Determination of native molecular weight of TRAPPIII and miniTRAPPIII by SEC-MALS. (E) GEF assays. Rabs were loaded with mant-GDP, and the change in fluorescence was measured over time after addition of a GEF or EDTA.

**Figure S2.**
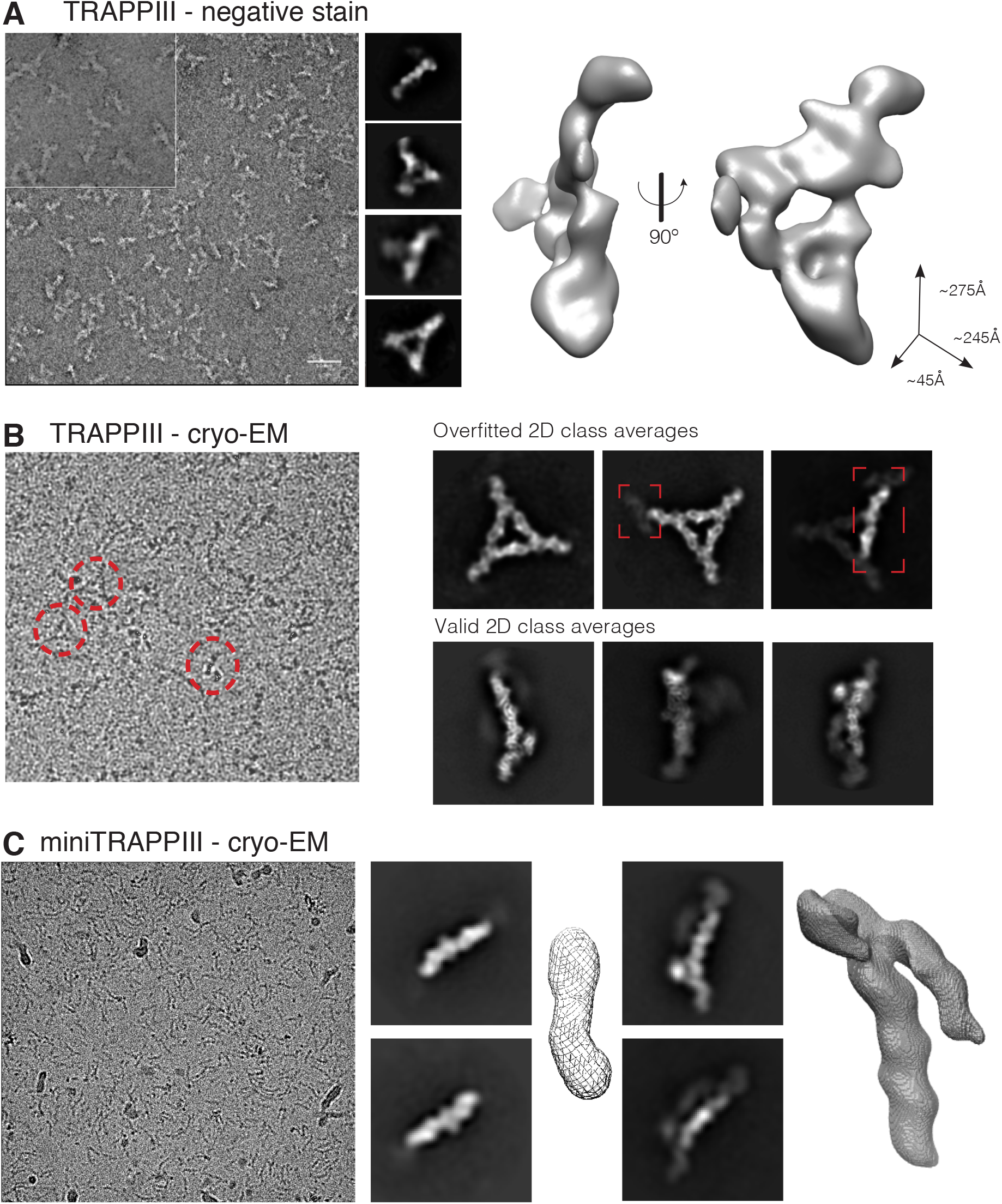
Negative staining and initial models. (A) Left: EM negative staining micrograph showing TRAPPIII particles. A zoomed inset with different contrast is shown in the top left corner of the micrograph. Scale bar 50 nm. Middle: Representative 2D classes. Right: 3D model from negative staining 2D classes. (B) Example cryo-EM micrograph of the whole TRAPPIII complex. The different particles shapes are highlighted with dashed circles. Overfitted and real 2D class averages are shown. (C) Example cryo-EM micrograph of miniTRAPPIII. 2D class averages and 3D initial models are shown.

**Figure S3.**
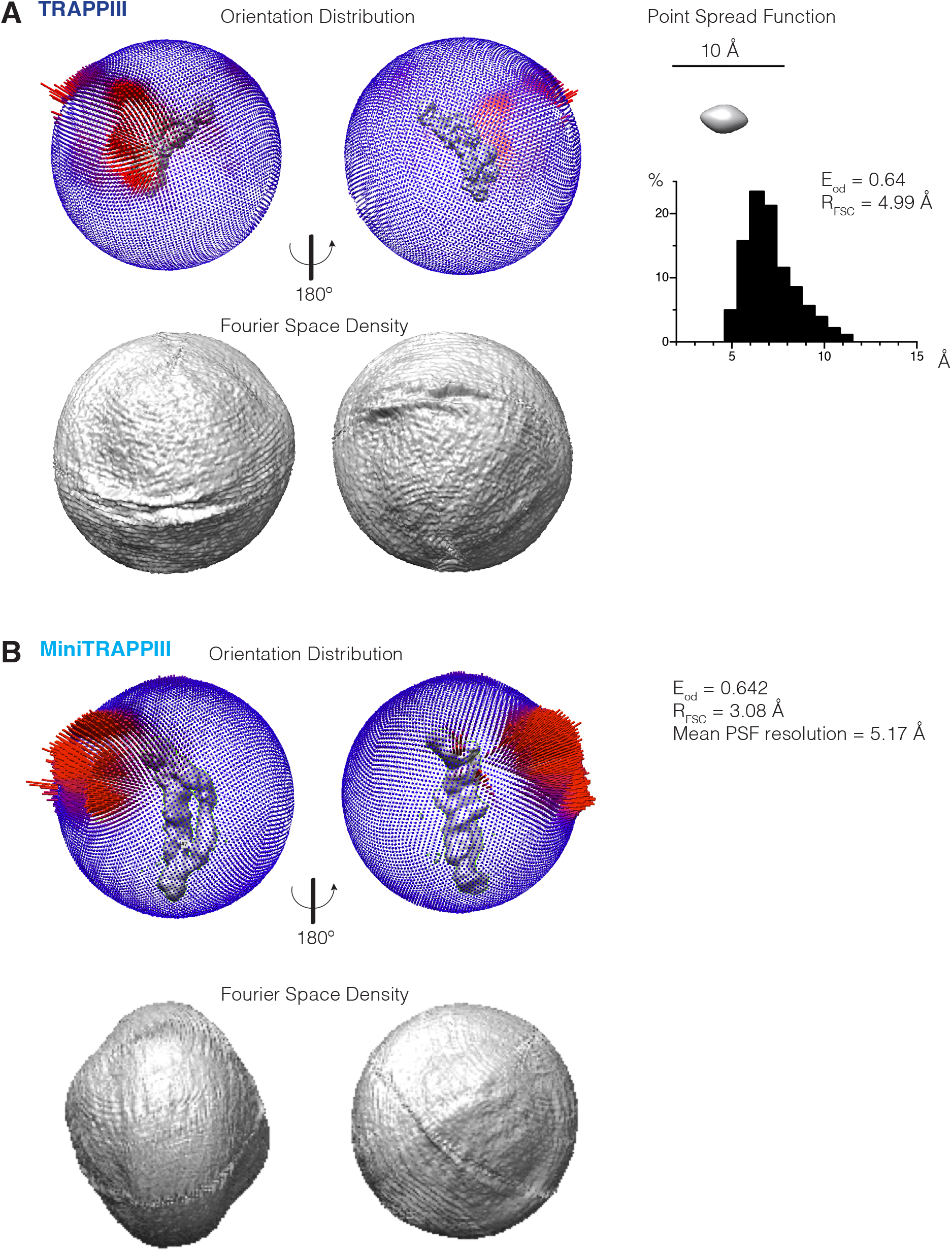
Angular distribution of non-tilted TRAPPIII and miniTRAPPIII. (A) and (B). Particle orientation distributions for TRAPPIII and miniTRAPPIII quantified by calculating their efficiency, *E*_od_ (Naydenova and Russo, 2017).

**Figure S4.**
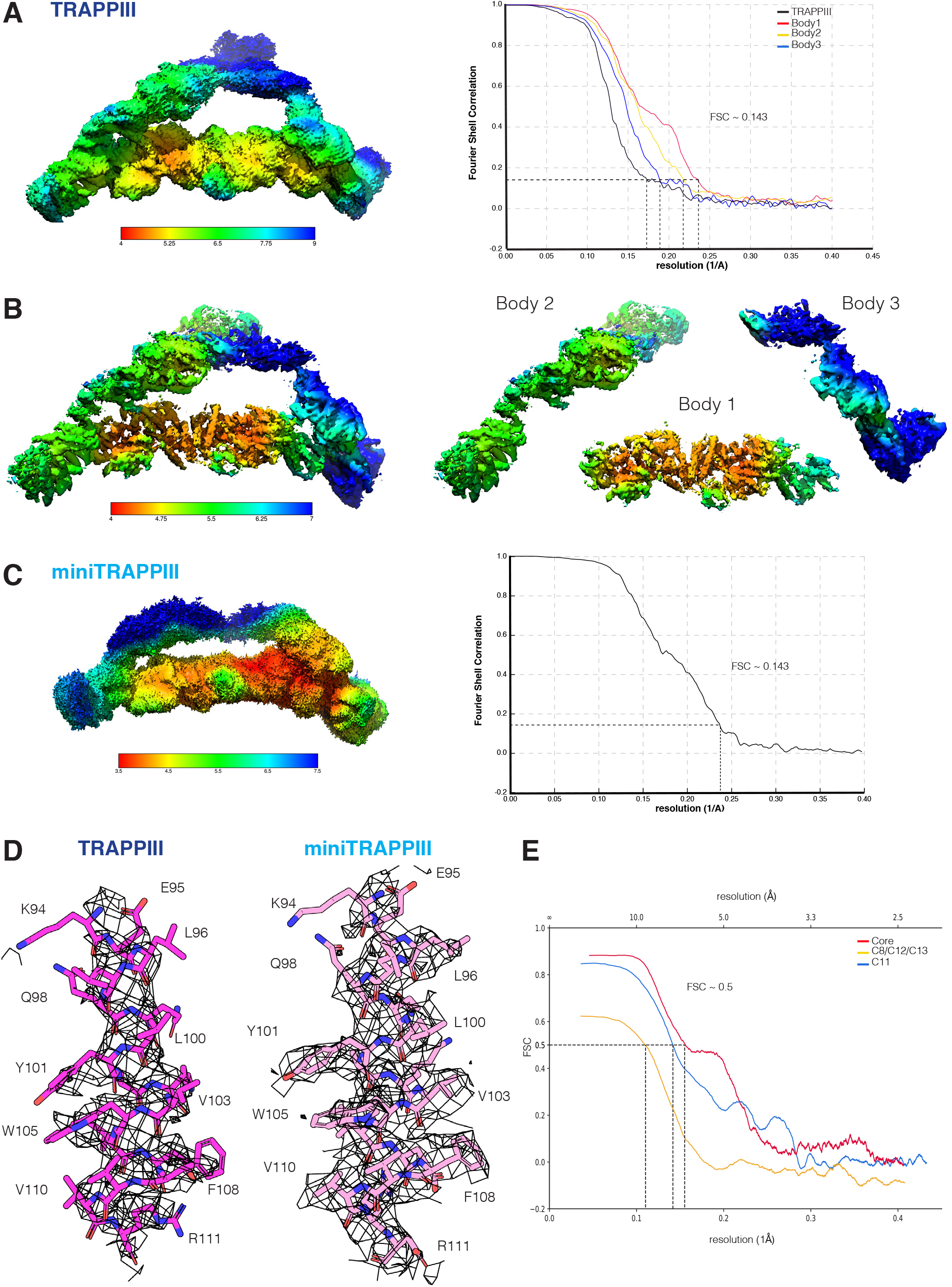
Cryo-EM structure of TRAPPIII and miniTRAPPIII. (A) TRAPPIII particles from two non-tilted and two tilted data sets were merged and 3D refinement was performed with RELION 3.1 to yield consensus maps at a nominal resolution of 5.8 Å. The consensus map from RELION was subjected to multi-body refinement, resulting in a major improvement. Shown is the local resolution estimation of the resulting cryo-EM map, and FSC curves indicating the overall nominal resolutions using the FSC = 0.143 criterion (body1/core ~ 4.2 Å, body2/C8-C12-C13 ~ 4.5 Å, body3/C11 −5.4 Å). (B) Local resolution estimation of the three different bodies used for multibody refinement. (C) MiniTRAPPIII particles from only one data set were refined with RELION 3.1 to yield consensus map at nominal resolution of 4 Å. The local estimation of the cryo-EM map and the FSC curve are shown. (D) Cryo-EM densities and refined models for representative regions of TRAPPIII and miniTRAPPIII. (E) FSC validation of the model versus cryo-EM map. The models refined against the first half-map (work), and the latter models versus the second half-map (free). The intersections of the curves with FSC=0.5 are shown.

**Figure S5.**
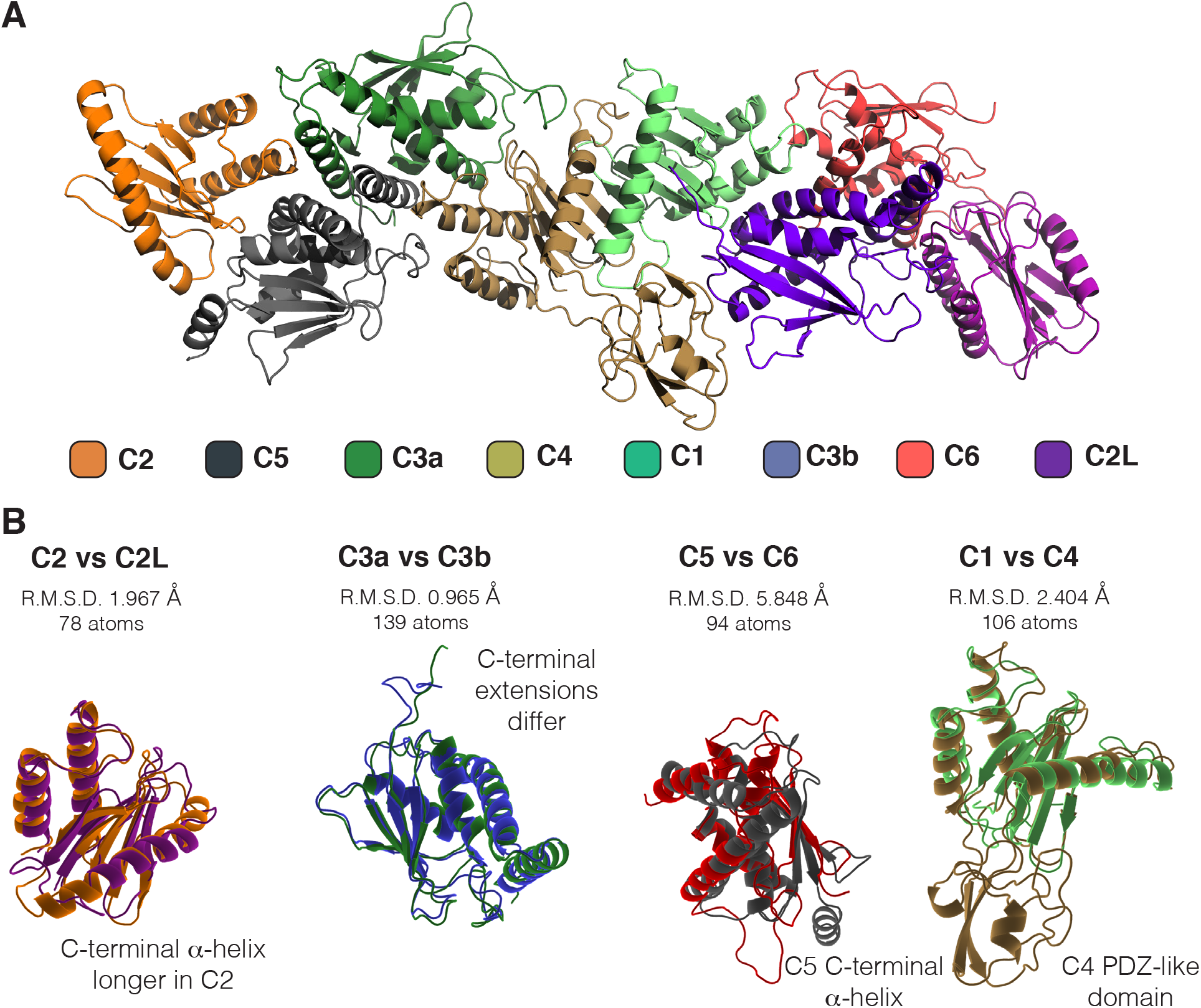
Arrangement of TRAPP core subunits. (A) Model of the core as fitted into the density map, with the eight subunits indicated by colour. (B) Comparison of the related pairs of subunits indicating their close similarity, with labelling to describe the additional features present in only one of the pair.

**Figure S6.**
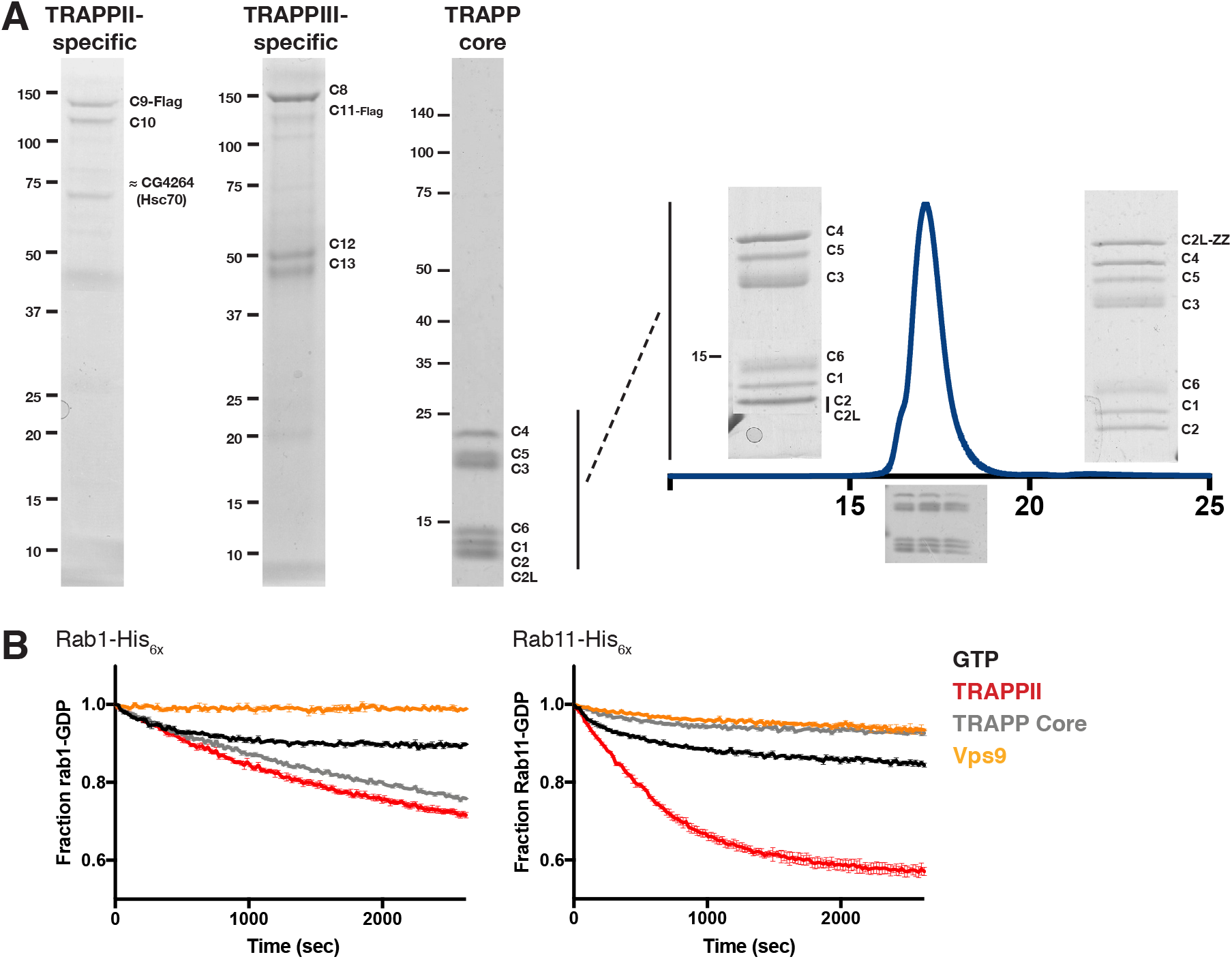
TRAPP subcomplexes. Coomassie blue–stained protein gels of purified TRAPP subcomplexes. FLAG tags on TRAPPC10 or TRAPPC11 allowed isolation of TRAPPII- or TRAPPIII-specific subunits, respectively, in the absence of core subunits. The core of TRAPP was purified by tagging TRAPPC3 with a Strep-Tag or TRAPPC2L with ZZ-domain. HRV-3C protease was used to cleave the tags before SEC. Normalized UV trace from SEC is shown. GEF assays performed in triplicate. Rabs were loaded with mant-GDP, and the change in fluorescence was measured over time after addition of a GEF. The Rab5 GEF Vps9 was used as a negative control.

**Figure S7.**
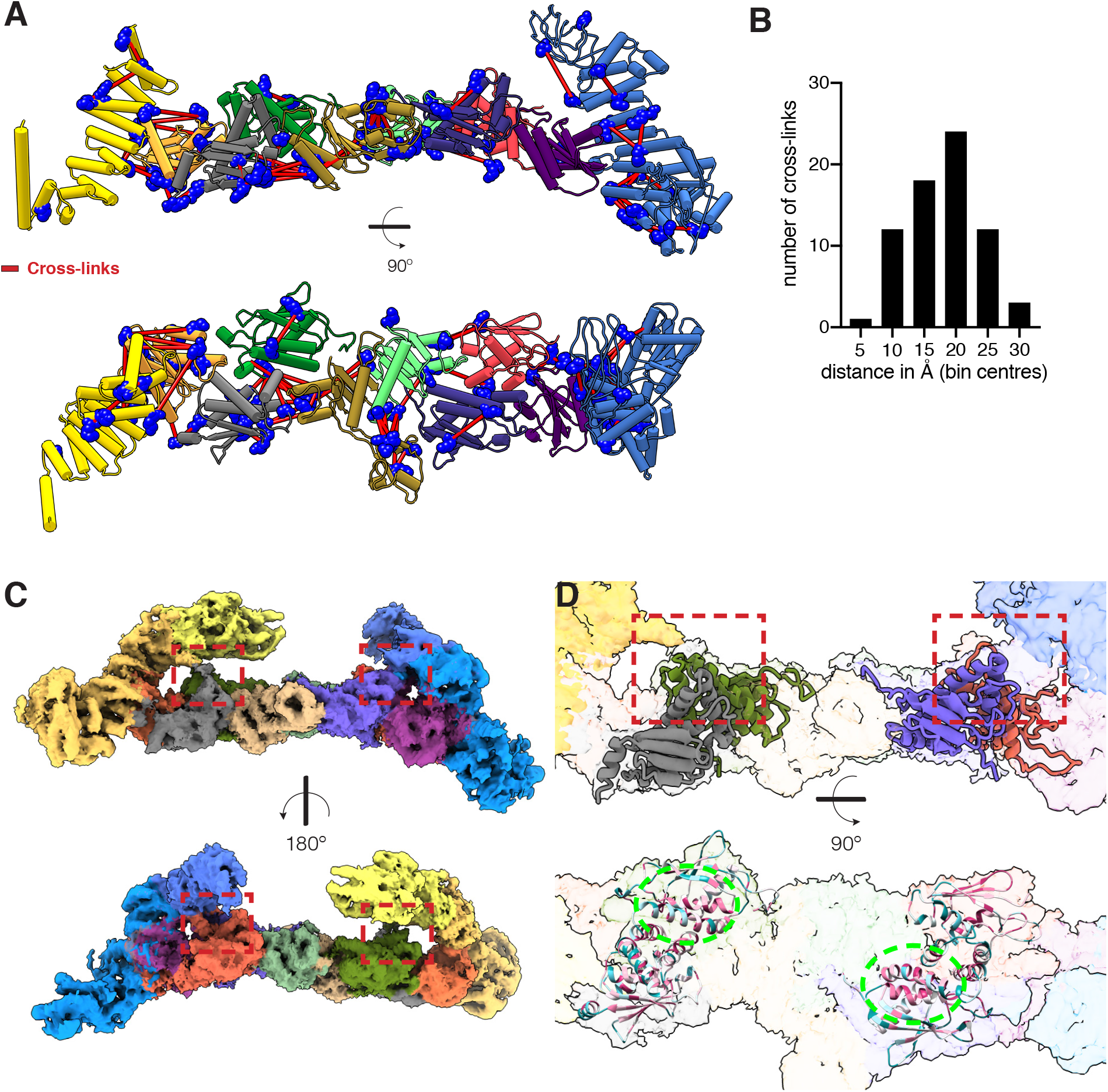
Cross-links and the TRAPPIII structural model. (A) Cross-links detected by mass-spectrometry mapped onto the structural model. All the detected cross-links falling within the expected Cα-Cα maximum distance threshold for DSBU are shown as red lines. The subunits are depicted as ribbon diagrams and differentiated by colour (C1: light green, C2: orange, C2L: magenta, C3a: purple, C3b: dark green, C4: light brown, C5: grey, C6: red, C8: yellow, C11: blue). Lysines in the structures are indicated as blue dots. (B) Bar graph shows the Cα-Cα distance distribution of all DSBU cross-links in the structure (maximum possible distance = ~ 30 Å). (C) Two views of the TRAPPIII density map coloured according to different subunits (C1: light green, C2: orange, C2L: magenta, C3a: purple, C3b: dark green, C4: light brown, C5: grey, C6: red, C8: yellow, C11: blue). The distal parts of the TRAPPC8 and TRAPPC11 arms have been removed for clarity. The density linking the core with TRAPPC8 and TRAPPC11 is indicated with a red dashed square. (D) Zoom of (C) showing the structural models of TRAPPC3a and TRAPPC6, and TRAPPC3b and TRAPPC5 fitted into the density near the bridges between the core and the arms. Subunits coloured as in (C). In bottom panel the subunits are coloured according to evolutionary conservation (red, most and blue, least). The part of the core that contacts the corresponding arm is formed by an arrangement of four αhelices that is partially conserved (green dashed oval).

**Figure S8.** Flexibility of TRAPPIII: Analysis of the relative orientations of the core and the TRAPPC8 and TRAPPC11 arms. (A) Results of the principal component analysis derived from the multibody refinement (Nakane et al., 2018). Inspecting the movement of the TRAPPIII complex relative to a body defined by the core and the N-terminal regions of TRAPPC8 and TRAPPC11 attached to it, a single eigenvector accounts for almost half of the variance. The histogram shows a unimodal distribution of the amplitude along this eigenvector, indicating a continuous range of positions. (B, C) Motion represented by the main eigenvector from multi-body analysis in (A). The vector represents a rocking motion of the core region in relation to the rest of the complex. This results in a rotation of the arms relative to the core of 13°, with a distance of around 18 Å between the two extreme positions of the part of TRAPPC8 that is predicted to touch Rab1 when the latter is bound to the catalytic site.

**Figure S9.**
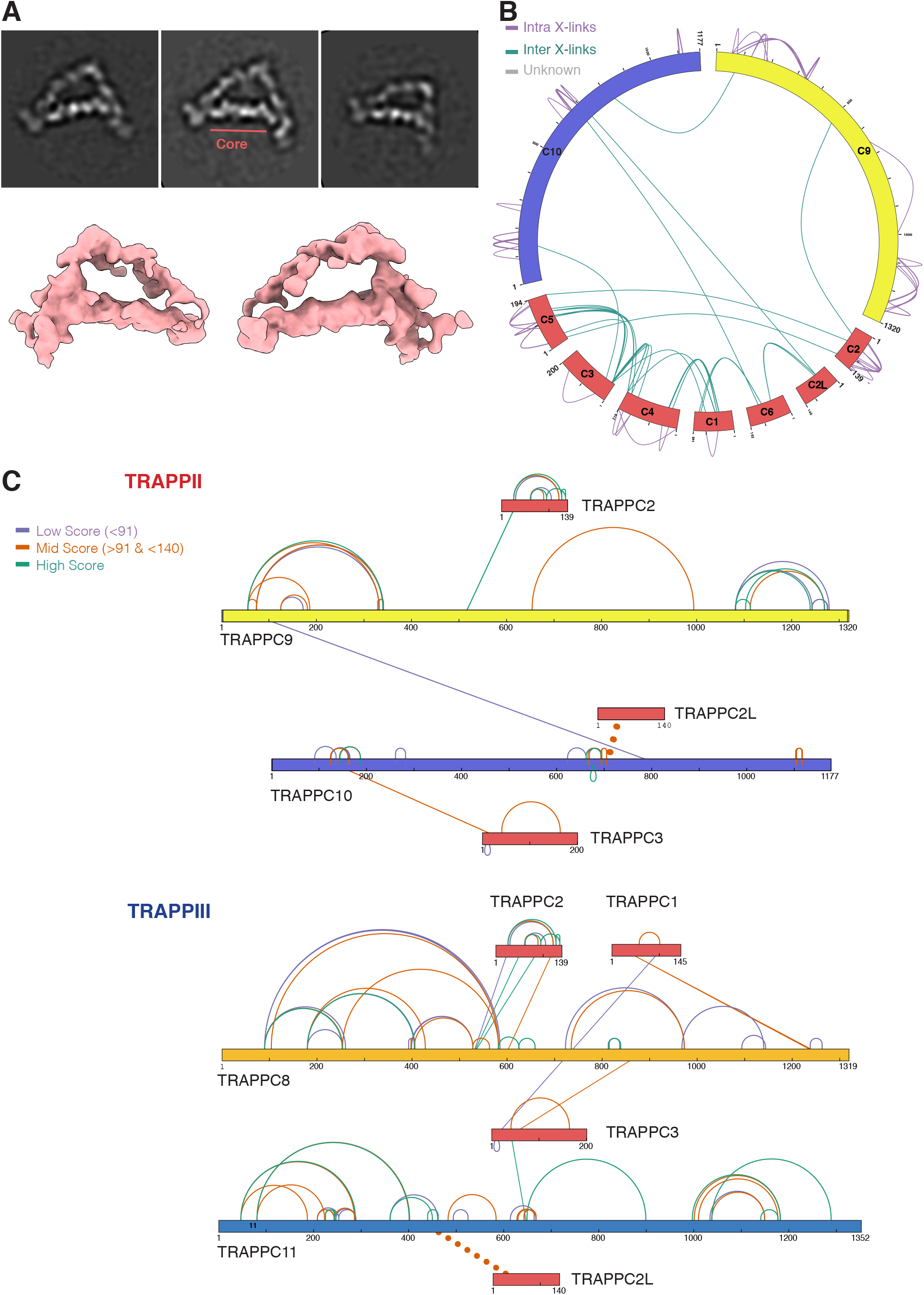
Architecture of the *Drosophila* TRAPPII complex. (A) Top: representative 2D class averages of TRAPPII. The core region is indicated. Bottom: low resolution cryo-EM map of TRAPPII. Circos-XL plot showing the distribution of all DSBU cross-links for the whole TRAPPII complex. Each protein is represented as a coloured segment (core subunits, red; TRAPPC9, yellow; TRAPPC10, purple), with residues numbers indicated. The relative position of the cross-link represents its location within the primary sequence. Inter-molecular cross-links are depicted as purples lines on the outside of the plot and intra-molecular cross-links as green lines inside of the plot. Network maps depicting the crosslinks found for TRAPPC9 (light yellow) and TRAPPC10 (purple) in TRAPPII, and TRAPPC8 (dark yellow) and TRAPPC11 (blue) in TRAPPIII. The cross-links are coloured according to the MeroX Score. The TRAPPC2 and TRAPPC2L cross-links are highlighted by dotted lines.

## Supplemental Information

**Table S1.** Crosslinking mass-spectrometry data for TRAPPII and TRAPPIII as well as the residues distances for the cross-links mapped on the TRAPPIII structural model.

**Video S1.** Movement of the arms relative to the core as defined by the principle vector of flexibility identified by multibody refinement.

